# A GSK3-dependent phosphorylation switch licenses BNIP3-mediated mitophagy

**DOI:** 10.64898/2026.07.24.740504

**Authors:** Anastasia V. Minenkova, Riya R. Philip, Yun Li, Thomas R. Hurd

## Abstract

The turnover of mitochondria through mitophagy is essential for maintaining mitochondrial function and matching mitochondrial content to cellular demand. In many cells, this process is mediated by the paralogous mitochondrial receptors BNIP3 and BNIP3L, which recruit WIPI-family autophagy effectors to initiate mitophagosome formation. However, the mechanisms that activate BNIP3/L remain poorly understood. Here, we show that the kinase GSK3 directly phosphorylates BNIP3, licensing receptor activity and activating mitophagy. Phosphorylation promotes BNIP3 recruitment of WIPI proteins, thereby initiating mitophagosome formation. We identify the critical phosphorylation sites required for this regulation and demonstrate that disruption of these sites abolishes BNIP3-dependent mitophagy. Together, our findings identify phosphorylation as a molecular switch controlling BNIP3/L activity and suggest that receptor phosphorylation may temporally and spatially license mitophagosome formation and mitochondrial clearance.

## INTRODUCTION

Mitophagy plays a central role in mitochondrial network remodeling by eliminating mitochondrial material to control organelle mass and maintain organelle integrity and cellular function (Montava-Garriga & Ganley, 2020; Vargas *et al*, 2023). Among the pathways that mediate mitophagy, the paralogous receptors BNIP3 and BNIP3L (also known as NIX) are increasingly recognized as key drivers of developmental and homeostatic mitophagy (Ney, 2015; Niemi & Friedman, 2024). These receptors reside on the outer mitochondrial membrane and promote phagophore formation and encapsulation of mitochondrial material for degradation via the endolysosomal system (Ney, 2015; Niemi & Friedman, 2024). Notably, BNIP3/L-driven mitophagy often occurs in a compartmentalized manner, with discrete subregions of the mitochondrial network selectively eliminated while adjacent regions are preserved (Yamashita *et al*, 2016; Niemi & Friedman, 2024). Yet BNIP3/L are broadly distributed across the mitochondrial surface, suggesting that additional mechanisms must regulate receptor activation during mitophagy initiation, potentially contributing to the spatial restriction of mitophagy within mitochondrial networks (Gok *et al*, 2023; Niemi & Friedman, 2024). However, the mechanisms that activate BNIP3/L on the mitochondrial surface remain poorly understood, despite substantial progress in defining pathways that regulate overall receptor abundance.

Global BNIP3/L levels are regulated at multiple levels, including transcriptional induction and post-transcriptional turnover. Transcriptional programs induce BNIP3/L expression under conditions such as hypoxia or during erythrocyte differentiation (Guo *et al*, 2001; Bruick, 2000; Sowter *et al*, 2001; Schweers *et al*, 2007; Sandoval *et al*, 2008), while multiple post-transcriptional pathways regulate global receptor turnover (Niemi & Friedman, 2024). Among these, the F-box protein FBXL4 ubiquitinates BNIP3/L and targets the receptors for proteasomal degradation in a process facilitated by the PP2C phosphatase PPTC7 (Niemi *et al*, 2023; Nguyen-Dien *et al*, 2024; Sun *et al*, 2024; Xu *et al*, 2025; Niemi *et al*, 2019; Wei *et al*, 2024; Elcocks *et al*, 2023; Cao *et al*, 2023; Nguyen-Dien *et al*, 2023; Chen *et al*, 2023), while the ER membrane protein complex (EMC) promotes receptor turnover through an autophagy-independent lysosomal pathway (Delgado *et al*, 2024). In addition, multiple phosphorylation events regulate global receptor abundance and stability, likely by modulating these and related turnover mechanisms (He *et al*, 2022; da Silva Rosa *et al*, 2021; Poole *et al*, 2021; Niemi *et al*, 2023).

By contrast, far less is understood about the mechanisms that directly activate BNIP3/L once it is on the mitochondrial surface. BNIP3/L contain three conserved functional elements that represent potential regulatory nodes: a LC3-interacting region (LIR) that engages Atg8/LC3 family proteins to facilitate phagophore encapsulation; a minimal essential region (MER, also known as the SLiM) that recruits WIPI proteins to promote local phagophore biogenesis; and a transmembrane domain that anchors the receptors within the outer mitochondrial membrane (Niemi & Friedman, 2024; Ney, 2015; Taoka *et al*, 2025).

Among these elements, the LIR appears the least critical, as mutations in this motif have only modest effects on mitophagy *in vivo* (Zhang *et al*, 2012; Novak *et al*, 2010) (see Fig. 6F), although more measurable effects have been observed in *in vitro* contexts (Ordureau *et al*, 2021; Bunker *et al*, 2023). Regulation of this motif by phosphorylation has been proposed based on phospho-mimetic analyses showing that substitution of residues flanking the LIR enhances mitophagy (Rogov *et al*, 2017; Zhu *et al*, 2013). However, phosphorylation of these residues has not been observed *in vivo* and phospho-null mutations do not significantly impair mitophagy (Bunker *et al*, 2023; Rogov *et al*, 2017; Zhu *et al*, 2013), indicating that phosphorylation at these sites, if it occurs, is unlikely to constitute a major mechanism of receptor activation.

In contrast to the LIR, both the transmembrane domain and MER are essential for receptor activity (Niemi & Friedman, 2024; Ney, 2015) and therefore represent plausible regulatory nodes. BNIP3/L transmembrane dimerization is critical for receptor activation, but the signals governing this transition are unknown (Marinković *et al*, 2021; Hanna *et al*, 2012; Sulistijo *et al*, 2003). Likewise, the mechanisms regulating MER-dependent WIPI recruitment, the key step in BNIP3/L mitophagy (Bunker *et al*, 2023; Adriaenssens *et al*, 2025), remain undefined.

Here, by investigating a developmentally programmed mitophagy event that occurs during Drosophila oogenesis, we identify a GSK3-dependent phosphorylation switch that is essential for BNIP3/L activation. We show that GSK3 associates with mitochondria at the onset of mitophagy, where it directly phosphorylates BNIP3 on conserved residues. Alanine substitution of these residues abolishes mitophagy, whereas phospho-mimetic variants bypass the requirement for GSK3. Mechanistically, BNIP3 phosphorylation enhances its interaction with the Drosophila WIPI orthologs Atg18a and Atg18b, thereby promoting mitophagosome formation. Notably, we find that this GSK3-dependent regulatory mechanism is conserved in human cells. Together, our findings identify GSK3 as an essential upstream activator of BNIP3 and reveal a conserved phosphorylation-dependent licensing mechanism that may contribute to the spatial and temporal regulation of mitophagy initiation.

## RESULTS

We previously identified a developmentally programmed mitophagy event that is initiated at meiotic entry and is essential for mitochondrial quality control during egg production (Lieber *et al*, 2019; Palozzi *et al*, 2022). This mitophagy is driven by the sole Drosophila ortholog of mammalian *BNIP3* and *BNIP3L* (Palozzi *et al*, 2022) (Fig. 1A). To investigate the mechanism underlying this critical process, we generated a Drosophila strain with endogenously tagged BFP-BNIP3 together with mCherry-Atg8a (Hegedűs *et al*, 2016) to mark autophagic structures (Fig. 1B, S1A). This system enabled quantitative assessment of BNIP3-dependent mitophagy *in vivo* by measuring the total volume of BNIP3-positive foci that colocalize with Atg8a in live samples.

**Figure 1.**
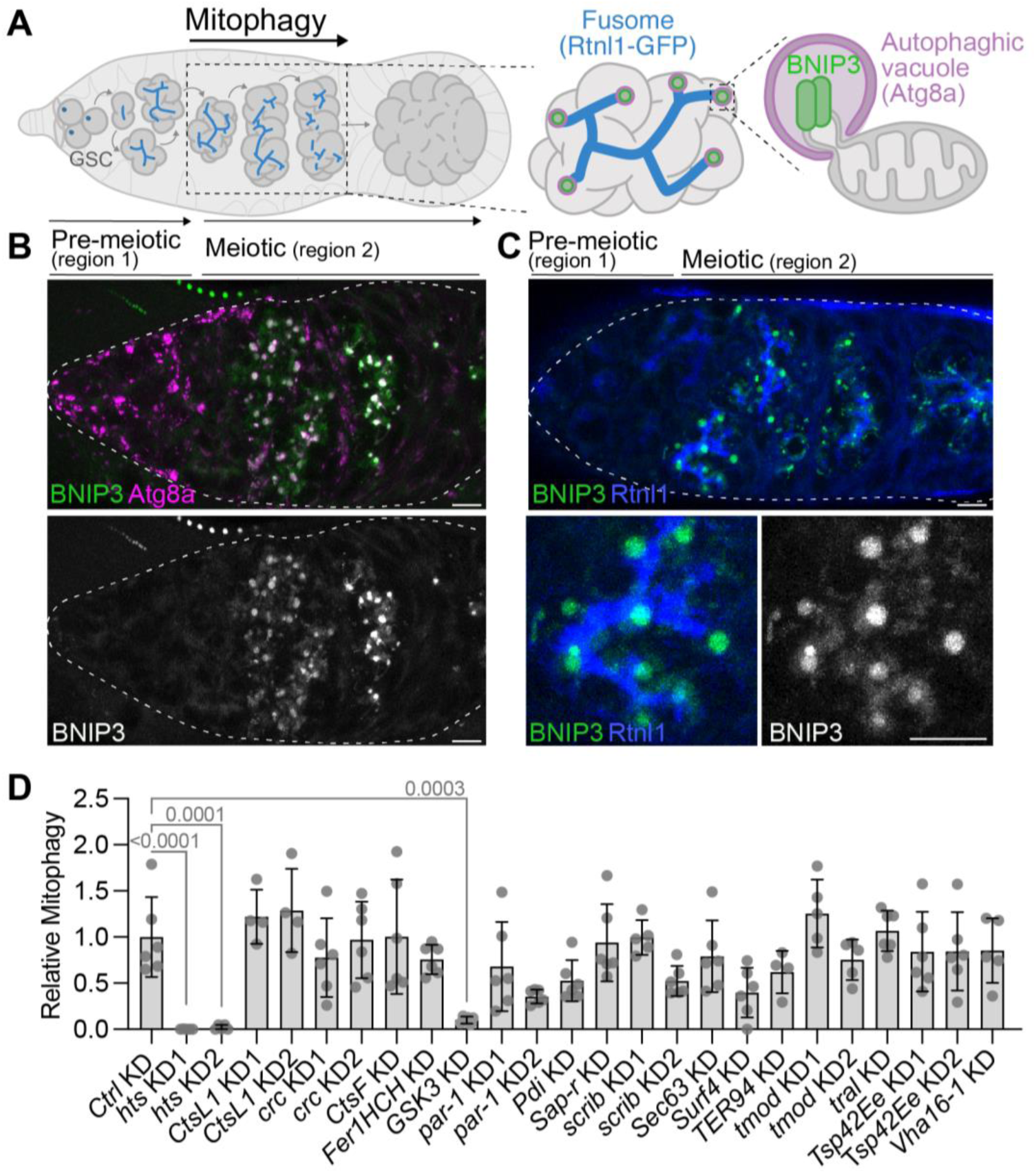
Screen identifies a role for GSK3 in germline mitophagy. **(A)** Schematic of *Drosophila* oogenesis. Germline stem cells (GSCs) at the anterior tip give rise to cells that undergo four mitotic divisions before entering meiosis. At meiotic onset, a robust BNIP3-dependent mitophagy program is activated and spatially organized by the fusome, an ER- and cytoskeleton-rich structure, marked by Rtnl1-GFP (blue). Mitophagosomes, marked by BFP-BNIP3 (green) and mCherry-Atg8a (magenta), associate with the fusome. **(B, C)** Confocal micrographs showing mitophagosomes labeled by BFP-BNIP3 (green) and mCherry-Atg8a (magenta) (**B**), or their association with the fusome (Rtnl1-GFP, blue) **(C)**. Max intensity projections are shown. White dashed lines outline the ovary. Scale bars, 5 µm. Genotypes: (**B**) *3xmCherry::Atg8a/+;BFP::BNIP3/+* and (**C**) *Rtnl1::GFP/+;BFP::BNIP3/+*. **(D)** RNAi screen identifies fusome-associated regulators of mitophagy. Knockdown of candidate genes in the germline (*nos-Gal4*) reveals a requirement for *hts* and *GSK3* in BNIP3-mediated mitophagy, quantified by overlap with Atg8a. Values are normalized to control KD (*LexA* shRNA on III) and represent means ± s.d.. *p*-values, one-way ANOVA with Dunnett’s correction. Only significant *p*-values (<0.05) are shown. Genotypes: progeny from *nos-Gal4 (NGT40); BFP::BNIP3, 3xmCherry::Atg8a x UAS-shRNA*.

In pre-meiotic germ cells, we observed low levels of BNIP3 that did not overlap with Atg8a, indicating that BNIP3 is largely inactive at this stage (Fig. 1B, S1B). Upon meiotic entry, however, BNIP3 protein levels increased significantly (Hsu & Drummond-Barbosa, 2017; Lieber *et al*, 2019), and we observed a pronounced increase of BNIP3 foci co-localizing with Atg8a, consistent with active receptor-mediated mitophagy (Fig. 1B, S1B). In addition to Atg8a, these BNIP3-positive foci also colocalized with endosomal (Rab7) and lysosomal (LAMP1 and lysotracker red) markers, further demonstrating that they represent mitochondria within autophagic vacuoles (hereinafter referred to as mitophagosomes) (Fig. S1C, C’). To confirm that BFP-BNIP3 is delivered to autophagic vacuoles in an autophagy-dependent manner, we knocked down the core autophagy regulator *Atg17*, which abolishes autophagy at these stages (Palozzi *et al*, 2022), and found that this likewise eliminated BNIP3 foci (Fig. S1D). Together, these data demonstrate that a robust BNIP3-mediated mitophagy program is initiated upon meiotic entry and establish BFP-BNIP3 as a faithful reporter of this receptor-mediated mitophagy.

### Screen for regulators of BNIP3-mediated germline mitophagy

Leveraging this developmentally programmed mitophagy as a physiologically relevant *in vivo* system, we sought to identify upstream activators of BNIP3. During the initial imaging, we observed that mitophagosomes were consistently associated with the fusome (Palozzi *et al*, 2022), a germline-specific ER–cytoskeletal organelle that interconnects developing germ cells (Huynh & St Johnston, 2004; de Cuevas *et al*, 1997) (Fig. 1A, C). Strikingly, BNIP3-positive mitophagosomes were frequently positioned at the termini of fusome branches, typically one per cell (Palozzi *et al*, 2022), suggesting that the fusome spatially organizes germline mitophagy upon meiotic entry (Fig. 1C). Consistent with this model, knockdown of *Hu-li tai shao* (*Hts*) (Yue & Spradling, 1992; Ding *et al*, 1993), a core structural component of the fusome, significantly reduced the abundance of BNIP3-positive mitophagic foci (Fig. 1D, S2). Together, these findings indicate that the fusome, or factors enriched at this structure, serves as a spatial platform for germline mitophagy and provides an entry point for identifying new BNIP3 regulators.

To identify fusome-associated factors required for this process, we performed a targeted RNAi screen of 17 genes (26 shRNAs) encoding proteins known to localize to the fusome (Lighthouse *et al*, 2008) and quantified mitophagy as described above. This screen identified a single additional factor, *shaggy* (Simpson *et al*, 1988), the sole Drosophila ortholog of *glycogen synthase kinase 3* (*GSK3*) *alpha* and *beta*, whose knockdown significantly impaired germline mitophagy (Fig. 1D). Assessment of fusome integrity by Hts immunofluorescence revealed no structural defects upon *GSK3* depletion or loss (Fig. S3A, B), indicating that the mitophagy defect is not secondary to fusome disruption. Furthermore, *GSK3* depletion did not decrease total BNIP3 protein levels (Fig. S3C), suggesting that GSK3 regulates BNIP3 activity rather than its abundance. Together, these findings identify GSK3 as a candidate activator of BNIP3-mediated mitophagy.

### GSK3 is required for BNIP3-mediated mitophagy

To determine whether GSK3 functions specifically in BNIP3-mediated mitophagy or instead plays a broader role in autophagy (Pan & Valapala, 2022), we quantified BNIP3- and Atg8a-positive foci in control and *GSK3* knockdown ovaries. If GSK3 were broadly required for autophagy in this context, its depletion would be expected to reduce both BNIP3 and Atg8a foci. Instead, while *GSK3* knockdown caused a marked reduction in BNIP3 foci (Fig. 2A, A′, S4), it had no effect on the volume of Atg8a foci (Fig. 2A, A″, S4), indicating that general autophagy remains intact and suggesting that GSK3 acts specifically in BNIP3-mediated mitophagy.

**Figure 2.**
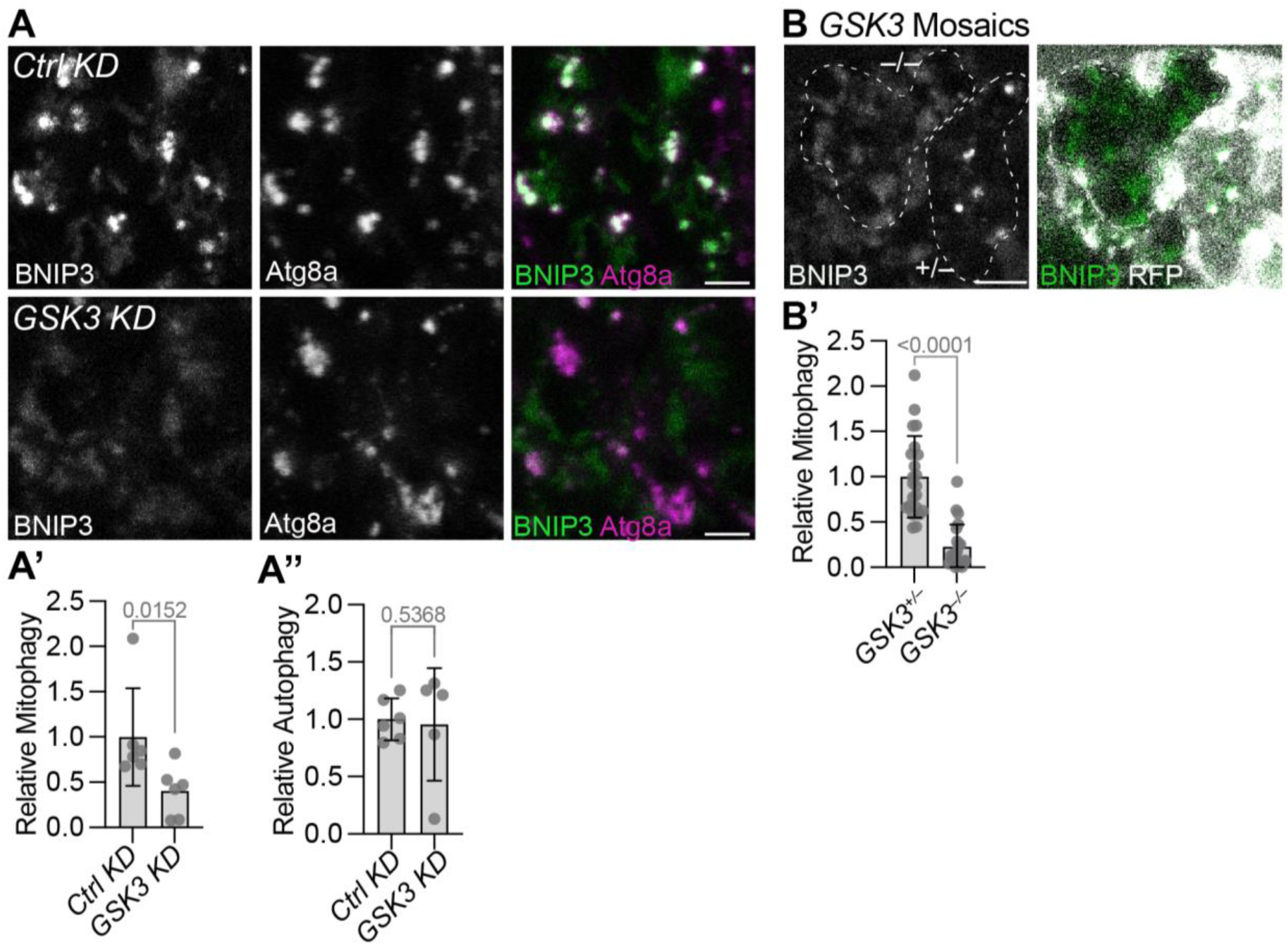
GSK3 as required for BNIP3-mediated germline mitophagy. **(A)** Zoomed-in confocal micrographs of BFP-BNIP3 and mCherry-Atg8a in control and *GSK3* germline (*nos*-Gal4) knockdown ovaries. Max intensity projections are shown. See Fig. S4 for full images. Scale bars, 2.5 µm. Genotypes: *Ctrl KD* = *NGT40/+;BFP::BNIP3, 3xmCherry::Atg8a/UAS-Lex shRNA | GSK3 KD* = *NGT40/+;BFP::BNIP3, 3xmCherry::Atg8a/UAS-GSK3 shRNA*. **(A’, A’’)** Quantification of the total volume of BFP-BNIP3 (A’) and mCherry-Atg8a (A’’) foci in (A). Values are normalized to control KD and represent means ± s.d. *p*-values, Mann-Whitney test. **(B)** Immunofluorescence confocal micrograph of BFP-BNIP3 in mosaic ovaries with *GSK3* mutant cells marked by loss of RFP. Max intensity projections are shown. White dashed lines outline the germline cysts. Scale bar, 2.5 µm. Genotype: *Ubi-mRFP, hsFLP, FRT19A/ GSK3^1^, FRT19A;;BFP::BNIP3/+*. **(B’)** Quantification of total volume of BNIP3 foci in heterozygote and homozygote *GSK3* mutant cells. Values are normalized to heterozygous control cells and represent means ± s.d. *p*-value, Mann-Whitney test.

To validate this phenotype, we confirmed that GSK3 was effectively silenced in our knockdowns by both RT-qPCR (Fig. S5A) and imaging (Fig. S5B). To exclude off-target effects of the RNAi, we assessed mitophagy in *GSK3* mutant germ cells using FLP/FRT-mediated mosaic analysis (Golic, 1991). Consistent with the knockdown phenotype, *GSK3* null cells (RFP-negative) exhibited a marked impairment in mitophagy relative to neighboring control cells (RFP-positive) (Fig. 2B, B′). Together, these data establish GSK3 as a specific regulator of BNIP3-mediated mitophagy in the Drosophila germline.

### GSK3 co-localizes with BNIP3 during mitophagy

To further dissect GSK3’s role in germline mitophagy, we examined its subcellular localization across the developmental transition into meiosis. We co-imaged BFP-BNIP3 with a GSK3-GFP protein trap strain (Lee *et al*, 2018; Wang *et al*, 2019), which is predicted to report the localization of all GSK3 isoforms detectably expressed in the ovary (>1 transcripts per million reads (Monteiro *et al*, 2026)). In pre-meiotic germ cells, GSK3 localized to the fusome, consistent with previous reports (Fig. 3A, arrows) (Lighthouse *et al*, 2008). Upon meiotic entry, however, GSK3 co-localized with BNIP3 in autophagic vacuoles (Fig. 3Ai), after which the GSK3-GFP signal was lost from the vacuoles (Fig. 3Aii), consistent with acidification of the autophagic compartment quenching the GFP fluorescence (Shinoda *et al*, 2018). Indeed, inhibition of acidification with chloroquine (Mauthe *et al*, 2018) prevented this signal loss, resulting in GSK3-GFP persisting within BNIP3-positive autophagic vacuoles (Fig. 3Aiii, iv) and increasing the colocalization between BNIP3 and GSK3 (Fig. 3A’).

**Figure 3.**
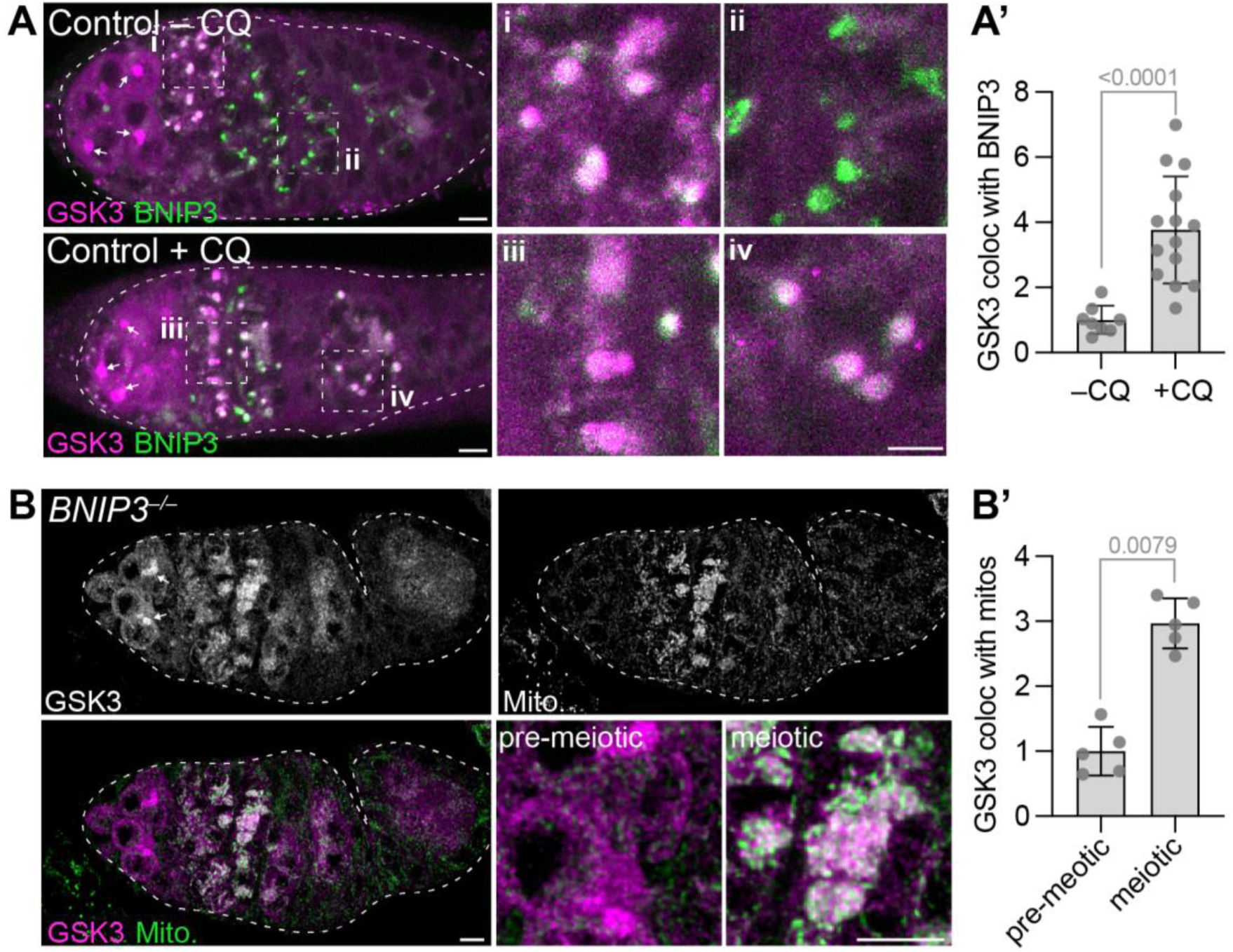
GSK3 associates with mitochondria independently of BNIP3. **(A)** Confocal micrographs of GSK3-GFP and BFP-BNIP3 in ovaries from flies fed with water (-CQ) or with chloroquine (+CQ). Arrows denote fusome associated GSK3-GFP. Max intensity projections are shown. White dashed lines outline ovaries. While dashed boxes indicate zoomed-in regions. Scale bars, 5 µm (whole) and 2.5 µm (zoom-ins). Genotype: *GSK3::GFP/+;;BFP::BNIP3/BFP::BNIP3*. **(A’)** Quantification of the colocalization of GSK3 with BNIP3 in mitophagosomes from (A). Values are normalized to the average of -CQ and represent means ± s.d.; *p*-value, Mann-Whitney test. **(B)** Confocal micrograph of *BNIP3* mutant ovaries expressing GSK3-GFP stained with anti-ATP5a (mito.) to mark mitochondria. Images were deconvolved using LAS X Lightning software (low signal-to-noise setting), and maximum-intensity projections are shown. White dashed line outlines the ovary. Scale bars, 5 µm and 2.5 µm (zoom-ins). Genotype: *GSK3::GFP/+;;BNIP3^1^/BNIP3^2^*. **(B’)** Quantification of colocalization of GSK3 with mitochondria in pre-meiotic vs meiotic stages from (B). Values are normalized to the average of pre-meiotic and are shown as means ± s.d. *p*-value, Mann-Whitney test.

We next asked whether GSK3 is recruited to mitochondria before mitophagosome formation or instead co-localizes with BNIP3 only after mitochondria have been engulfed. To distinguish between these possibilities, we blocked mitophagy by mutating *BNIP3* (Fig. S6) and examined GSK3 localization. In the absence of BNIP3-dependent mitochondrial engulfment, GSK3 accumulated on mitochondria upon meiotic entry, where mitochondria progressively accumulated because they were no longer be degraded (Fig. 3B). Thus, GSK3 recruitment to mitochondria occurs independently of autophagic engulfment. Moreover, mitochondrial recruitment of GSK3 did not require BNIP3 itself, indicating that GSK3 is targeted to the mitochondrial surface through a BNIP3-independent mechanism. Quantification confirmed a significant increase in GSK3-mitochondria colocalization upon meiotic entry in *BNIP3* mutants relative to pre-meiotic germ cells (Fig. 3B′). Together, these findings indicate that GSK3 is recruited to mitochondria at the onset of meiosis, where it promotes BNIP3-mediated mitophagy.

### GSK3 promotes mitophagy by activating BNIP3 through phosphorylation

Given that GSK3 is a kinase, we next asked whether it promotes mitophagy by phosphorylating a target protein, focusing on BNIP3 as a prime candidate. Analysis of phospho-proteomics data from iProteinDB (Hu *et al*, 2019) identified ten phosphorylated residues in Drosophila BNIP3 (Fig. 4A, BNIP3^WT^). None of these sites lie within known functional motifs (Niemi & Friedman, 2024; Ney, 2015).

**Figure 4.**
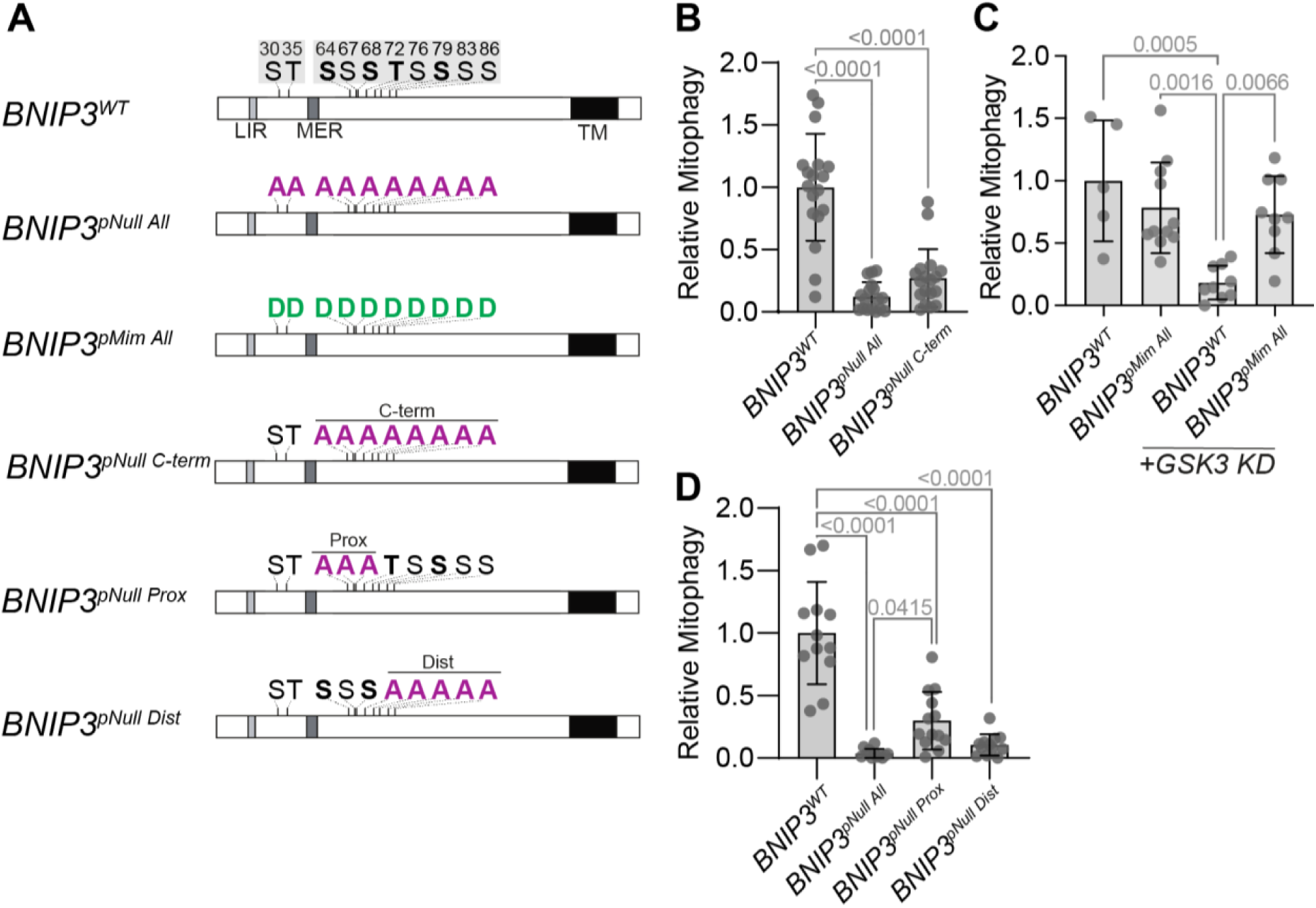
GSK3 promotes the phosphorylation and activation of BNIP3. **(A)** Schematic of wild-type and mutant BNIP3 showing potential phosphorylated residues, which cluster into two regions, N-terminal and C-terminal (grey boxes). Sites predicted to be phosphorylated by GSK3 are indicated in bold. The four predicted GSK3 phosphorylation sites localize to the C-terminal block, which can be further subdivided into proximal and distal regions. **(B, C, D)** Quantification of mitophagy in control and *BNIP3* mutant backgrounds. BNIP3-mediated mitophagy was quantified by overlap with Atg8a (**B**) or by measuring total amount of BFP-BNIP3 foci (**C**, **D**). See Figs. S7, S8 and S9 for images. Values are normalized to *BNIP3^WT^* and shown as means ± s.d. *p*-values, one-way ANOVA with Tukey’s correction. Only significant *p*-values (<0.05) are shown. Genotypes: (**B**) *BNIP3^WT^* = *3xmCherry::Atg8a/+; BFP-BNIP3^WT^/BNIP3^1^ | BNIP3^pNull^ ^All^ = 3xmCherry::Atg8a/+; BFP-BNIP3^10S/T>A^/BNIP3^1^ | BNIP3^pNull^ ^C-term^* = *3xmCherry::Atg8a/+; BFP-BNIP3^8S/T>A^/BNIP3^1^.* (**C**) BNIP3^WT^ = *BFP::BNIP3^WT^; BNIP3^1^/BNIP3^2^ | BNIP3^pMim^ ^All^* = *BFP::BNIP3^10S/T>D^; BNIP3^1^/BNIP3^2^ | BNIP3^WT^ + GSK3 KD* = *NGT40/BFP::BNIP3^WT^; BNIP3^1^/ BNIP3^2^, UAS-GSK3 shRNA | BNIP3^pMim^ All + GSK3 KD* = *NGT40/BFP::BNIP3^10S/T>D^; BNIP3^1^/BNIP3^2^, UAS-GSK3 shRNA*. (**D**) *BNIP3^WT^ = BFP::BNIP3^WT^; BNIP3^1^/BNIP3^2^ | BNIP3^pNull^ ^All^* = *BFP::BNIP3^10S/T>A^; BNIP3^1^/BNIP3^2^ | BNIP3^pNull^ ^Prox^* = *BFP::BNIP3^3S/T>A^; BNIP3^1^/BNIP3^2^ | BNIP3^pNull^ ^Dist^* = *BFP::BNIP3^5S/T>D^; BNIP3^1^/BNIP3^2^*.

To determine whether these residues are required for BNIP3 function, we generated CRISPR-edited flies carrying alanine substitutions at all ten sites while retaining the endogenous BFP tag to assess mitophagy (Fig. 4A, BNIP3^pNull^ ^All^). Despite normal BNIP3 protein stability (Fig. S7A), germline mitophagy was severely impaired in the *BNIP3* phospho-null mutant (Fig. 4B, S7C) and was indistinguishable from those observed in either *GSK3* null mutants (p=0.9819) (Fig. S7B) or a completely non-functional *BNIP3* mutant lacking both its essential functional motifs (LIR and MER, p=0.9905) (Fig. 6F). These results indicate that phosphorylation of one or more of these residues is essential for BNIP3 function.

If GSK3 promotes mitophagy through phosphorylation of BNIP3, then mimicking phosphorylation at these sites should bypass the requirement for GSK3. Because the generation of additional CRISPR alleles proved technically challenging, we instead expressed a phospho-mimetic *BFP-BNIP3* variant, in which all ten sites were mutated to aspartic acid, under endogenous regulatory control in a *BNIP3* mutant background (Fig. 4A, BNIP3^pMim^ ^All^). We generated an otherwise identical wild-type transgenic strain in parallel as a control. Owing to the genetic complexity of these strains, we were unable to introduce the mCherry-Atg8a reporter and therefore quantified total BFP-BNIP3 foci, rather than their overlap with Atg8a, as a surrogate measure of mitophagy. This approach is justified by our earlier finding that >95% of BFP-BNIP3 foci co-localize with Atg8a and other markers of autophagic vacuoles (Fig. S1C’). We found that mitophagy occurred at levels comparable to controls in flies expressing the phospho-mimetic protein, indicating that the mutant receptor is functional (Fig. 4C, S8). Strikingly, *GSK3* knockdown no longer impaired mitophagy in the phospho-mimetic background, in contrast to flies expressing wild-type BNIP3 (Fig. 4C, S8). Together, these results indicate that GSK3 acts upstream of BNIP3 to promote its phosphorylation and activation during germline mitophagy.

We next examined whether any of these phosphorylation sites matched GSK3 consensus motifs. The sites clustered into two regions: an N-terminal block containing two sites and a C-terminal block containing eight sites (Fig. 4A). Notably, four sites within the C-terminal block match the canonical GSK3 consensus motif (S/T)XXX(S/T), in which GSK3 phosphorylates the C-terminal serine/threonine residue (Fiol *et al*, 1987, 1988) (Fig. 4A, BNIP3^WT^ bold). To assess the functional importance of the C-terminal block, we mutated all eight sites using CRISPR (Fig. 4A, BNIP3^pNull^ ^C-term^), as kinases can exhibit promiscuous phosphorylation of adjacent residues when primary sites are disrupted (Johnson *et al*, 2023). Mutation of the C-terminal eight sites impaired mitophagy to the same extent as mutation of all ten sites (p=0.3006) (Fig. 4B, S7C), indicating that the C-terminal sites account for the essential phosphorylation-dependent activity.

We next subdivided these sites into two groups, each with two predicted GSK3 sites, and generated transgenic mutants targeting either the three proximal sites or the five distal ones (Fig. 4A, BNIP3^pNull^ ^Prox^ and BNIP3^pNull^ ^Dist^). Mutation of the three proximal sites caused a partial reduction in mitophagy, whereas mutation of the five distal sites impaired mitophagy to a similar extent as mutation of all ten sites (p=0.8970) (Fig. 4D, S9). Together, these results indicate that phosphorylation of one or more C-terminal, distal residues is essential for BNIP3 function, while the proximal sites make a more modest contribution.

### GSK3 directly phosphorylates BNIP3 on serine 79

We next asked whether GSK3 directly phosphorylates BNIP3 *in vitro* and, if so, which of the eight key candidate residues are targeted. Using an ADP-Glo kinase assay, which measures ADP production as a proxy for kinase activity, we found that recombinant human GSK3 robustly phosphorylates purified BNIP3 lacking its transmembrane domain, which was removed to improve solubility (Fig. 5A, S10A to C). Mutation of all ten phosphorylation sites, all eight C-terminal phosphorylation sites, or either the proximal or distal C-terminal cluster alone reduced phosphorylation, suggesting that GSK3 directly phosphorylates BNIP3 at sites in both the C-terminal proximal and distal regions (Fig. 5A), consistent with our *in vivo* data.

**Figure 5.**
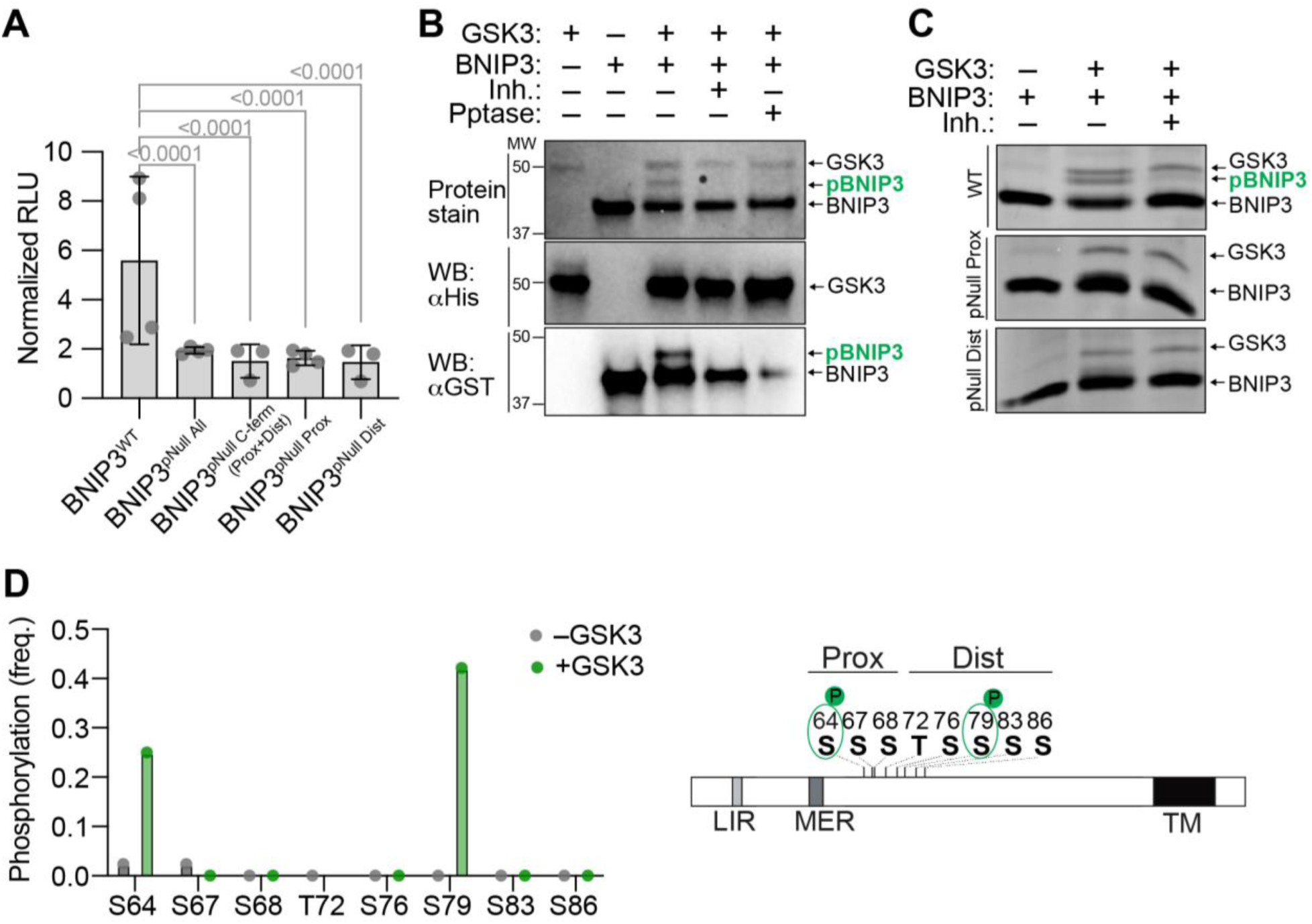
GSK3 directly phosphorylates BNIP3. **(A)** *In vitro* kinase activity as measured by relative luminescence unit (RLU) using ADP-Glo kit. GST-tagged wildtype and phospho-null BNIP3 mutants were incubated with or without human His-GSK3B. Values are normalized to the activity measured in the absence of the kinase (paired for each substrate). See Fig. S10. Each dot represents independent experimental mean of technical triplicates. *p*-values, one-way ANOVA with Tukey’s correction. Only significant *p*-values (<0.05) are shown. **(B)** Protein-stained polyacrylamide gel and western blots showing BNIP3 phosphorylation as measured by mobility shift. Gels were stained with InstantBlue Coomassie (top panel) and western blots with either anti-His (middle panel) or anti-GST (bottom panel). GST-BNIP3 was incubated with or without human His-GSK3B, in the prescence or absence of GSK3-specific inhibitor, CHIR99021 (Inh.) or λ-phosphatase (Pptase). **(C)** Protein-stained polyacrylamide gel showing BNIP3 phosphorylation as measured by mobility shift in BNIP3 bands. Gels were stained with InstantBlue Coomassie. GST-tagged variants of BNIP3, including BNIP3^WT^ (top panel), BNIP3^pNull^ ^Prox^ (middle panel) and BNIP3^pNull^ ^Dist^ (bottom panel), were incubated with or without His-hGKS3. **(D)** Phosphorylation frequency at individual BNIP3 residues following incubation with or without GSK3, as determined by LC–MS/MS analysis of *in vitro* phosphorylated BNIP3.

To directly assess phosphorylation, we turned to electrophoretic mobility shift assays. Incubation of BNIP3 with GSK3 produced a clear mobility shift that was blocked by the GSK3 inhibitor CHIR-99021 (Ring *et al*, 2003) and reversed by phosphatase treatment, confirming that the shift reflects phosphorylation (Fig. 5B). Consistent with the ADP-Glo results, mutation of the three C-terminal proximal or the five distal sites abolished the shift (Fig. 5C). This further demonstrates that both proximal and distal residues in C-terminal block are required for phosphorylation *in vitro*.

To precisely map which sites GSK3 phosphorylated, we performed mass spectrometry of BNIP3 following incubation with or without GSK3. Peptides spanning all ten candidate phosphorylation sites were recovered at varying abundance; however, phosphorylation was detected at only two of the predicted GSK3 target residues following addition of GSK3 (Fig. 5D, S10D, Table S2). Quantification of phosphorylated versus total peptide spectral counts at each residue revealed modest phosphorylation at serine 64 within the proximal C-terminal cluster and robust phosphorylation at serine 79 within the distal C-terminal cluster. Together, these data show that GSK3 directly phosphorylates BNIP3, with serine 79 representing the primary target site likely required for its mitophagy function.

### GSK3-mediated phosphorylation of BNIP3 promotes binding to Atg18/WIPI2

How GSK3-dependent phosphorylation activates BNIP3 remains unclear. Mitophagy receptors function by recruiting autophagy machinery to the mitochondrial surface, and recent work indicates that BNIP3 promotes turnover primarily through recruitment of WIPI family proteins (known as Atg18 in *Drosophila*) (Bunker *et al*, 2023; Adriaenssens *et al*, 2025). We therefore asked whether phosphorylation alters the interaction profile of BNIP3 to promote recruitment of downstream effectors.

To address this, we immunoprecipitated full-length phospho-null and phospho-mimetic BNIP3 from Drosophila S2 cells and performed mass spectrometry to identify proteins enriched with the activated phospho-mimetic form (Fig. 6A). Spectral counts from three independent replicates were averaged and normalized to untransfected controls, and proteins enriched ≥2-fold in the phospho-mimetic vs the phospho-mutant immunoprecipitation with a Bayesian false discovery rate (BFDR) ≤ 0.01 were considered significant interactors (Fig. 6B, Table S3). To prioritize physiologically relevant candidates, we filtered these hits using our prior RNA-seq dataset (Monteiro *et al*, 2026) for those transcripts enriched in early meiotic germ cells, where germline mitophagy occurs (Fig. 6C).

**Figure 6.**
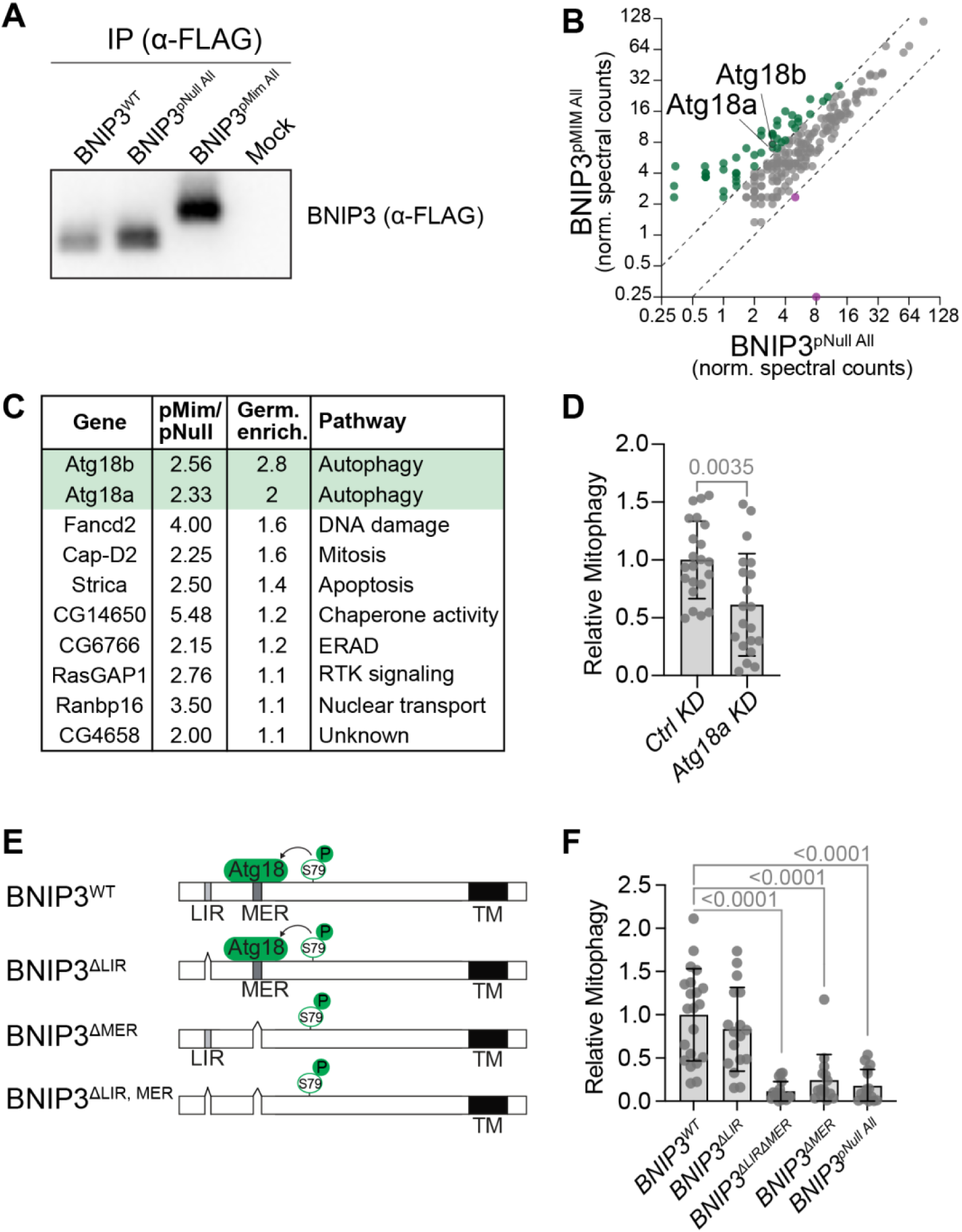
GSK3-mediated BNIP3 phosphorylation promotes recruitment of WIPI family proteins, Atg18a and b. **(A)** Representative western blot of wildtype, phospho-null and phospho-mimetic BNIP3 co-immunopreciptated from Drosophila S2 cells. Mock, untransfected cells. **(B)** A scatterplot showing the normalized spectral counts in phospho-mimic and phospho-null BNIP3. Spectral counts are averaged across three replicates. Dashed lines show 2-fold difference between the conditions. Shown are interactors significantly enriched in the pulldowns relative to the negative control with BFDR ≤ 0.01. **(C)** Table of top ten interactors enriched in phospho-mimetic BNIP3 immunoprecipitation condition relative to the phospho-null, as ordered by enrichment in meiotic germ cells relative to whole ovaries. **(D)** Knockdown of *Atg18a* inhibits mitophagy *in vivo* as quantified by the overlap of BNIP3 with Atg8a. Values are normalized to control knockdown and shown as mean ± s.d.; *p*-value, Mann-Whitney test. See Fig. S11 for images. Genotypes: *Ctrl KD* = *NGT40/UAS-LexA shRNA; 3xmCherry::Atg8a, BFP::BNIP3/+ | Atg18a KD* = *NGT40/UAS-Atg18a shRNA; 3xmCherry::Atg8a, BFP::BNIP3/+*. **(E)** A cartoon showing the structure-function mutants of BNIP3 generated with CRISPR. **(F)** Quantification of mitophagy in ovaries expressing wild-type *BNIP3* (*BNIP3^WT^*), *BNIP3* lacking LIR (*BNIP3^ΔLIR^*), *BNIP3* lacking MER (*BNIP3^ΔMER^*), BNIP3 lacking both LIR and MER (*BNIP3^ΔLIRΔMER^*) or phospho-null *BNIP3*. Mitophagy is quantified by the overlap between BNIP3 and Atg8a. Values are normalized to BNIP3^WT^ and shown as means ± s.d. *p*-values, one-way ANOVA with Tukey’s correction. Only significant *p*-values (<0.05) are shown. See Fig. S13 for images. Genotypes: *BNIP3^WT^*= *3xmCherry::Atg8a/+; BFP-BNIP3^WT^/BNIP3^1^ | BNIP3^ΔLIR^*= *3xmCherry::Atg8a/+; BFP-BNIP3^ΔLIR^/BNIP3^1^ | BNIP3^ΔLIRΔMER^* = *3xmCherry::Atg8a/+; BFP-BNIP3^ΔLIRΔMER^/BNIP3^1^ | BNIP3^ΔMER^* = *3xmCherry::Atg8a/+; BFP-BNIP3^ΔMER^/BNIP3^1^ | BNIP3^pNull^* = *3xmCherry::Atg8a/+; BFP-BNIP3^p10S/T>A^/BNIP3^1^*.

This analysis identified the WIPI orthologs Atg18a and Atg18b as top candidates (Fig. 6C, Table S3). Indeed, consistent with a functional role, depletion of *Atg18a*, the more highly expressed of the two paralogs in the female germline (Monteiro *et al*, 2026), impaired germline mitophagy *in vivo* (Fig. 6D, S11). As expected for a core WIPI family protein, *Atg18a* depletion also reduced the number of Atg8a-positive autophagic foci, consistent with an additional role in autophagy (Shimizu *et al*, 2023). When we performed a similar analysis comparing wild-type and phospho-mutant immunoprecipitations, we again detected an interaction with Atg18b (Fig. 6A, S12, Table S4). Interestingly, we also detected an interaction between BNIP3 and GSK3 itself, consistent with direct association of the kinase with its substrate, although this interaction is not required for GSK3 localization to the mitochondrial surface, as GSK3 remained mitochondrial in the absence of *BNIP3* (Fig. 3C). Together, these data support a model in which GSK3-mediated phosphorylation of BNIP3 promotes Atg18 binding to facilitate mitophagy.

We next sought to define how phosphorylation promotes WIPI recruitment. BNIP3 contains both a LIR, which canonically binds LC3 family proteins (Zhang *et al*, 2012; Novak *et al*, 2010), and a MER, which mediates WIPI recruitment (Bunker *et al*, 2023; Adriaenssens *et al*, 2025). However, recent work has also implicated the LIR in WIPI binding (Adriaenssens *et al*, 2025). To determine whether phosphorylation promotes WIPI recruitment through the LIR or MER, we disrupted each domain individually (Fig. 6E). If phosphorylation acts primarily through the LIR, deletion of this motif should phenocopy the phospho-null mutant; conversely, if phosphorylation acts through the MER, disruption of the MER should produce a similar defect. Strikingly, deletion of the LIR had only a modest effect on mitophagy, whereas deletion of the MER impaired mitophagy to the same extent as the phospho-null mutant (p=0.9887, Fig. 6F, S13A). Neither the LIR nor the MER mutation affected protein stability (Fig. S13B). Together, these data indicate that GSK3-dependent phosphorylation does not activate BNIP3 exclusively through the LIR and suggest that it may instead promote MER-dependent WIPI recruitment to enable germline mitophagy.

### GSK3-mediated BNIP3 activation promotes mitophagy in human cells

Lastly, we wanted to explore if human BNIP3/L is regulated by GSK3 in a similar fashion to Drosophila BNIP3. The key serine residue S79 phosphorylated in Drosophila BNIP3 is conserved in human BNIP3/L (Fig. 7A). We monitored mitophagy in HeLa cells expressing the mito-QC reporter following deferiprone (DFP) treatment (Zhao *et al*, 2020), which robustly induces BNIP3-dependent mitophagy (Fig. 7B, C). Strikingly, pharmacological inhibition of GSK3 with CHIR-99021 abolished DFP-induced mitophagy. These findings suggest that GSK3 activity is also required for BNIP3/L-dependent mitophagy in mammalian cells and support a conserved mechanism of BNIP3 activation.

**Figure 7.**
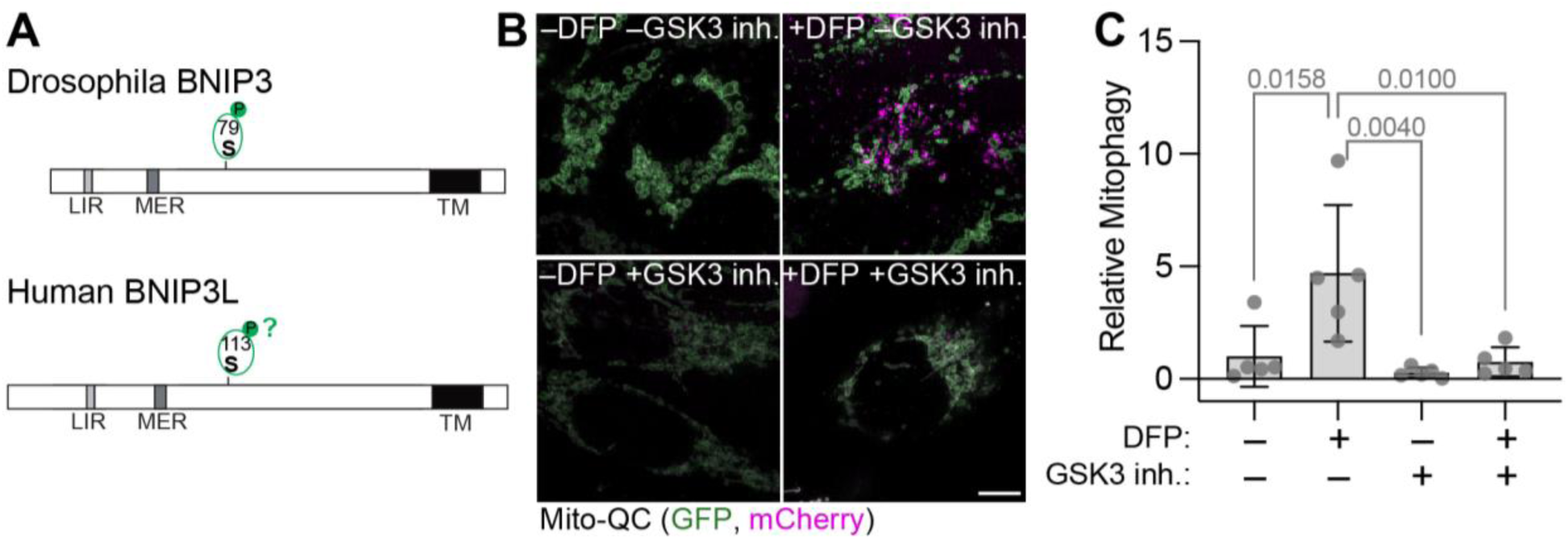
GSK3-mediated BNIP3 phosphorylation promotes mitophagy in human cells. **(A)** Schematic showing the relative position of the key GSK3-phosphorylated residue in Drosophila BNIP3 and the corresponding residue in human BNIP3L. **(B)** Confocal micrographs of HeLa cells transiently expressing a mitophagy reporter mito-QC while being treated with or without deferiprone (DFP) and GSK3 inhibitor (CHIR-99021). Maximum intensity projections of are shown. Scale bar, 5 µm. **(C)** Quantification of mitophagy in HeLa cells. Values are normalized to the untreated cells (no DFP, no CHIR) and shown as means ± s.d.. Each value (dot) represents the data collected from ∼100 cells. *p*-values were calculated using one-way ANOVA with Tukey’s correction. Only significant *p*-values are shown.

## DISCUSSION

Here, we identify a conserved GSK3-dependent phosphorylation switch that activates BNIP3/L-mediated mitophagy. We show that phosphorylation of BNIP3 promotes WIPI recruitment and is essential for mitophagy in both Drosophila and human cells, thereby establishing phosphorylation-dependent receptor licensing as a conserved mechanism regulating BNIP3/L activity. Our findings provide a framework for understanding how BNIP3/L-mediated mitophagy is regulated with spatial and temporal precision.

A major outstanding question is how phosphorylation mechanistically activates BNIP3/L. We identify serine 79 as a critical regulatory residue in Drosophila BNIP3 and find that phosphorylation of BNIP3 promotes WIPI recruitment. Because WIPI recruitment is sufficient to initiate local phagophore biogenesis (Adriaenssens *et al*, 2025), these findings place phosphorylation at one of the earliest committed steps of BNIP3-mediated mitophagy. This mechanism is distinct from previously proposed models involving phosphorylation of residues flanking the LIR, which primarily influence LC3 binding and have more modest effects on mitophagy (Bunker *et al*, 2023; Rogov *et al*, 2017; Zhu *et al*, 2013). Instead, our data indicate that GSK3-dependent phosphorylation activates BNIP3 by promoting WIPI recruitment, likely through the MER. How phosphorylation facilitates this interaction remains an important question. Phosphorylation may induce conformational changes that expose or stabilize the MER or alternatively generate a docking site for an as-yet unidentified factor that promotes WIPI recruitment. Distinguishing between these possibilities will be important for understanding how BNIP3/L transitions from an inactive mitochondrial resident to an active mitophagy receptor.

An equally important question is whether BNIP3/L phosphorylation is spatially regulated within mitochondrial networks. BNIP3/L-mediated mitophagy frequently occurs in a compartmentalized manner, with discrete regions of the mitochondrial network eliminated while adjacent regions are preserved (Yamashita *et al*, 2016; Niemi & Friedman, 2024). Localized GSK3-dependent phosphorylation provides one plausible mechanism for generating this spatial specificity. Consistent with this possibility, we find that GSK3 localizes to the mitochondrial surface independently of BNIP3 itself, suggesting that mitochondrial pools of GSK3 may be selectively positioned or activated at sites of mitophagy initiation. GSK3 localization is regulated by multiple scaffold proteins, including AKAP family proteins, several of which associate with mitochondria (Wu *et al*, 2022), making them attractive candidates for spatially restricting BNIP3/L activation. In addition, many GSK3 substrates require priming phosphorylation by upstream kinases (Patel & Woodgett, 2017), raising the possibility that BNIP3/L activation is controlled through combinatorial kinase signaling. Identification of the upstream priming kinase(s), the mechanisms that recruit or activate GSK3 at the mitochondrial surface, and the phosphatases that reverse BNIP3/L phosphorylation will therefore be important future directions.

Lastly, our findings have broader implications for understanding how developmentally programmed mitophagy is regulated and functions. In many developmental contexts, including germline mitochondrial quality control and erythrocyte maturation, mitophagy is initiated through upregulation of BNIP3/L protein (Schweers *et al*, 2007; Sandoval *et al*, 2008; Lieber *et al*, 2019; Palozzi *et al*, 2022). Our findings suggest that developmental induction of BNIP3/L establishes a window of competence for mitophagy, whereas GSK3-dependent phosphorylation provides a second layer of regulation that licenses receptor activity. Such a two-tiered mechanism could allow developmental programs to specify when mitophagy can occur while simultaneously permitting spatial restriction of receptor activation within mitochondrial networks. In the Drosophila germline, this additional layer of regulation may be particularly important for mitochondrial genome quality control, where selective elimination of dysfunctional mitochondrial subdomains, rather than wholesale mitochondrial turnover, is thought to be critical for purifying selection (Palozzi *et al*, 2022). It will therefore be important to determine whether GSK3-dependent receptor licensing similarly regulates other forms of developmentally programmed mitophagy and contributes more broadly to mitochondrial remodeling during development. Interestingly, GSK3 has also been implicated in mitochondrial remodeling and the induction of quiescence during later stages of *Drosophila* oogenesis (Yue *et al*, 2022; Sieber *et al*, 2016), raising the possibility that GSK3-dependent activation of BNIP3 contributes more broadly to developmental mitochondrial remodeling.

Collectively, our findings support a model in which receptor-mediated mitophagy is regulated not simply through receptor abundance but through local post-translational licensing of receptor activity at the mitochondrial surface. This mechanism provides a means to integrate developmental, metabolic, and spatial cues, allowing cells to regulate not only when mitophagy occurs but also where it occurs within continuous mitochondrial networks. More broadly, regulated receptor licensing may represent a general principle by which cells spatially and temporally control mitophagy while preserving organelle integrity.

## ACKNOWLEDGMENTS

We thank Chalini Weerasooriya and Brandon Payliss for help with optimising conditions for the *in vitro* kinase assays. We thank Toby Lieber, Mayu Shimomura and Yusuke Murase for helpful comments on the manuscript. We thank Bloomington Drosophila Stock Center (NIH P40 OD018537) for Drosophila stocks and Addgene for plasmids. The Hts (1B1) antibody (DSHB Cat# 1b1, RRID:AB_528070) was obtained from the Developmental Studies Hybridoma Bank (DSHB), created by the NICHD of the NIH and maintained at The University of Iowa, Department of Biology, Iowa City, IA 52242. We thank Dr. Gino Laberge from Genome Prolab for assitance with generating the transgenic lines. We thank Rainbow Transgenic Flies, Inc for generating the CRISPR lines. We thank Dr. Michael Moran, Dr. Craig Simpson and Owen Tsai at SPARC Biocentre, Hospital for Sick Children for assistance with the phospho-proteomics. We thank Cassandra Wong of the Network Biology Collaborative Centre Proteomics Facility (RRID: SCR_025375) at the Lunenfeld-Tanenbaum Research Institute for the IP/MS-MS in S2 cells. The facility is supported by the Canada Foundation for Innovation and the Ontario Government. This work was supported by the Canadian Institutes of Health Research and T.R.H. is part of the University of Toronto Medicine by Design initiative, which receives funding from CFREF.

## AUTHOR CONTRIBUTIONS

Conceptualization, R.R.P., A.V.M., and T.R.H.; methodology, R.R.P., A.V.M., and T.R.H.; reagents, R.R.P., A.V.M., and T.R.H.; investigation, R.R.P., and A.V.M.; writing – original draft, R.R.P., A.V.M., and T.R.H.; writing – review & editing, R.R.P., A.V.M., and T.R.H.; funding acquisition, T.R.H.; supervision, T.R.H and Y.L.

## DATA AVAILIBILTY STATEMENT

Strains generated in this study are available from the authors upon reasonable request. The authors affirm that all other data necessary for confirming the conclusions of the article are present within the article, figures, tables and supplementary materials.

## MATERIALS AND METHODS

### Drosophila Husbandry and Stocks

All fly stocks were reared on standard medium (cornmeal, yeast, agar, and molasses) at 25°C and 60-70% humidity on a 12-hour light/12-hour dark cycle. All knockdown and mutant experiments were conducted at 25 °C. All stocks used are listed in Table S1. We used FlyBase (release FB2025_05) (Öztürk-Çolak *et al*, 2024) to obtain stock, expression and sequence information.

### Generation of *GSK3 (sgg)* mosaics

The *sgg^1^* (also known as *sgg^b12^*) allele has an inversion breakpoint in the middle of the *GSK3* gene and has been wildly used as a genetic null (Takeo *et al*, 2012). *sgg^1^/FM7* (BDSC #4095) virgin females were crossed to *FRT19A* (BDSC #1744) males to generate recombinants. *sgg^1^, FRT19A/FM7* were crossed to *Ubi-mRFP, HsFLP, FRT19A/Y;;BFP::BNIP3/TM3Sb* males and 2^nd^ and 3^rd^ instar larvae were heat-shocked at 37°C twice for 2 hours each, with a recovery period of 4 hours in-between. *sgg^1^, FRT19A/Ubi-mRFP, HsFLP, FRT19A;;BFP::BNIP3/*+ female progeny were yeasted 7 days after heat-shock and imaged on the 8^th^ day. Germline cysts that were mRFP-negative were identified as *sgg^1^*-positive clones, which were compared to mRFP-positive cysts as internal negative controls.

### Generation of *BNIP3* strains using CRISPR/Cas9

#### BNIP3^1^ and BNIP3^2^ loss-of-function alleles

We generated two independent *BNIP3* mutant alleles, *BNIP3^1^*and *BNIP3^2^*, using CRISPR/Cas9-mediated genome editing (Gratz *et al*, 2015). Both alleles introduce frameshift mutations predicted to truncate BNIP3 before the critical MER and transmembrane (TM) domains. Consistent with this, *BNIP3* transcript was largely absent in *BNIP3^1/2^*transheterozygotes (Fig. S6).

*BNIP3^1^* was generated by injecting a pCFD4 plasmid (Addgene #49411; (Port *et al*, 2014)) expressing two gRNAs (5′-GGCAACGGAGTGATTCTGTC-3′ and 5′-GGGACTGTTCGGTGGACTCT-3′), assembled using the NEBuilder HiFi DNA Assembly Master Mix (NEB E2621L), into vasa-Cas9 embryos (BDSC #51324). Individual F0 adults were crossed to a balancer strain, and F1 progeny were used to establish independent lines. Genome editing was identified by PCR and confirmed by Sanger sequencing using the following primers: forward, 5′-ACGCTGAAAGGGTGACGATG-3′; reverse, 5′-AAGGCCGCAATGCTCCAAAC-3′. *BNIP3^1^* carries a frameshift at Pro31 predicted to introduce a premature stop codon at Lys46.

*BNIP3^2^* was generated during CRISPR/Cas9-mediated insertion of the endogenous BFP tag (see below). This allele carries a frameshift at Ala24 predicted to introduce a premature stop codon at Asp71.

#### Endogenous BFP tagging of BNIP3

We inserted a mTagBFP2 tag into the *BNIP3* locus using CRISPR/Cas9-mediated homologous recombination. A donor construct containing mTagBFP2, ∼1.25 kb left and ∼0.85 kb right homology arms, and BNIP3 genomic sequence (exon 2 to 5) harboring silent mutations within the gRNA target sites and linker regions was assembled in the Scarless-HD-DsRed-w+ vector (Addgene #80801; a gift from Kate O’Connor-Giles) (see Fig. S1A). mTagBFP2 was PCR-amplified from pCAG mito-mTagBFP2 (Addgene #105011, (Kwon *et al*, 2016)) and left and right homology arms (LHA and RHA respectively) were amplified from BL51324 flies using the following primers:

LHA fw = 5′-GCGTCTACCTGACATACG-3′
LHA rev = 5′-CTGGAAAGTAGATCATTAGCATG-3′
RHA fw = 5′-GCCATAAACAAAAAGGAGAAAGAGG-3′
RHA rev = 5′-GCAGCACTTCCCTGAAAATCC-3′.

BNIP3 fragments were synthesized by Twist Bioscience (Table S5). DsRed-positive F1 progeny were identified by eye fluorescence, precise integration was confirmed by Sanger sequencing, and the selectable marker was excised by crossing to a PiggyBac transposase line (BDSC #8285).

#### Generation of BNIP3 mutant alleles

*BNIP3 ΔMER, ΔLIR, ΔLIRΔMER*, *phospho-null (10S/T-A; pNull All),* and *C-terminal phospho-null (8S/T-A; pNull C-term)* alleles were synthesized by Twist Bioscience (Table S5) and introduced into the endogenous *BNIP3* locus using the same CRISPR/Cas9-mediated strategy described above.

### Generation of transgenic BNIP3 phospho-mutant strains

*pBNIP3-BFP-BNIP3* plasmids were assembled using PCR amplification with 2× GB-AMP PaCeR HP Master Mix (GeneBio Systems) and Gibson assembly using the NEBuilder HiFi DNA Assembly Master Mix (NEB, E2621) into pCasper (gift from Dr. Uli Tepass). We amplified the *BNIP3* regulatory regions (5’ and 3’ UTRs) from *w^1118^* genomic DNA using the following primers:

5’ UTR fw = 5′-ACTTTTCATATTCATATATGCATATG-3 ′,
5’UTR rev = 5′-CGCCCAGCAAATCTTCGC-3′,
3’UTR fw = 5′-AAGATGCCATAAACAAAAAG-3′,
3’UTR rev = 5′-ACACAAAGGTAATCCAAGCCC-3′.

DNA fragments encoding BFP fused to wild-type or mutant *BNIP3* were synthesized by Twist Bioscience and assembled into the expression vector. Transgenic flies were generated by ΦC31-mediated integration into the attP40 landing site (*y^1^, w^1^; P{y^[+t7.7]^=CaryP}attP40*) by Genome ProLab.

### Chloroquine Feeding

1-to-2-day old *GSK3-GFP* expressing flies were sorted and placed on low melt food (7% corn syrup (v/v), 1.5% agarose mixture [11:1 – low melt agarose: regular agarose] (w/v), 2% yeast (w/v), in distilled water) supplemented with 3mg/mL chloroquine (Bioshop CHL919) in distilled water. Flies were fed food with chloroquine or water for three days, with fresh food on the first and third day.

### Imaging Live Ovaries

All mitophagy and autophagy analysis was conducted on live ovaries. Ovaries from flies feed with yeast overnight were dissected in PBS and incubated in CellMask™ Deep Red Plasma Membrane Stain (1:1000 dilution, Invitrogen, Cat# C10046) for 2 minutes and rinsed with fresh PBS. Ovaries were then placed on a coverslip in Halocarbon 700 oil (Sigma-Aldrich H8898). Individual ovarioles were pulled out from the muscular sheath using Tungsten needles and positioned them on the coverslip such that all stages are expanded and spread out. A live imaging chamber that contains a gas permeable membrane was then placed over the coverslip containing the mounted ovaries, which was then imaged on a Leica SP8 inverted scanning confocal microscope using a 63X, NA 1.4 immersion oil objective.

### Immunostaining Fixed Ovaries

Adult ovaries were immunostained according to standard procedures. Briefly, ovaries from yeasted flies were dissected in PBS and fixed in 4% formaldehyde (Thermo Scientific, 28908) in PBS for 15 min. Ovaries were then permeabilized with 1% Triton X-100 (BioShop Canada, TRX506) in PBS for 20 min. Ovaries were incubated with primary antibodies (anti-Hts, 1:20; anti-ATP5a, 1:1000, Abcam ab14748) diluted in PBST (1% BSA BioShop Canada, ALB001, 0.1% Triton X-100, PBS) overnight at 4°C followed by incubation with the appropriate secondary antibodies (anti-mouse Cy3, 1:500, Jackson La) diluted in 1% PBST for 2 hours at room temperature. Ovaries were mounted in VECTASHIELD Antifade Mounting Medium (Chromatographic Specialties Inc. VECTH19002). All images were acquired with a Leica SP8 inverted scanning confocal microscope using 63x, NA 1.4, immersion oil objectives.

### Mitophagy quantification in ovaries

To quantify mitophagy events in the germline, fluorescent reporters BFP-BNIP3 and mCherry-Atg8a were used. Surfaces were generated in Imaris (v.9). Briefly, background was subtracted using local contrast threshold with the diameter of the largest sphere determined as the diameter of the largest object of interest in each image and was kept constant within each experiment (1-1.20µm). For BNIP3 surfaces, manual threshold value was established based on brightness to better capture concentrated BNIP3 within mitophagosomes and was kept the same within each experiment. For Atg8a surfaces, the automatic brightness threshold was used. Only surfaces within the germarium and that were greater than 10.0 voxels were kept. The sum of “Overlapped Volume to Surfaces” between BNIP3 and Atg8a surfaces was used and normalized to the germline volume, which was determined manually by outlining the germline using Cell Mask^TM^. In cases where the mCherry-Atg8a could not be included, mitophagy was quantified by summing the volume of BNIP3 surfaces alone since they colocalized with the degradative markers (Figure S1C-C’). For all analyses, the data are compared to matched control samples to account for differences in genetic background.

### Colocalization analysis

#### Colocalization between BNIP3 and degradative markers

For all colocalization analyses Imaris v.9 was used. BNIP3 surfaces were generated the same way as for mitophagy quantitation. LAMP1, Lysotracker, and Rab7 surfaces were generated similarly to how Atg8a surfaces were generated.

#### Colocalization between BNIP3 and GSK3

BNIP3 surfaces were generated using the same method as for mitophagy quantification with the following changes: intensity cutoff was set lower than for the mitophagy quantifications to account for the increased size of mitophagosomes, sphericity cutoff was set to >0.585 and volume cutoff was set to <7.4µm^3^. For GSK3 surfaces, background was subtracted using local contrast threshold with the diameter of the largest sphere set to 2.00µm. Manual threshold brightness cutoff was used to avoid capturing diffuse cytosolic GSK3 signal and was kept the same for all images. Only surfaces within the germarium and larger than 10.0 voxels were kept. Colocalized volume of BNIP3 and GSK3 was determined from the sum of “Overlapped Volume to Surfaces” and normalized to the germline volume.

#### Colocalization between GSK3 and mitochondria

Surfaces for pre-meiotic and meiotic regions were generated manually using the changes in morphology of the germline developmental stages as a guidance. Mitochondrial surfaces were generated from ATP5a signal. Background was subtracted using the local contrast threshold with surface grain size of 0.1µm and diameter of largest sphere of 1.8µm. Automatic brightness threshold was used. Split touching Objects function was enabled with Seed Points Diameter = 0.4µm. Lower “Quality” objects were eliminated using automatic quality cutoff. Only the surfaces within the germarium and larger than 10.0 voxels were kept. GSK3 surfaces were generated using the same method except the diameter of largest sphere of 2.00µm was used. Colocalized volume of ATP5a and GSK3 was determined from the sum of “Overlapped Volume to Surfaces” and normalized to the volume of the corresponding region.

### BNIP3 intensity quantification in the ovaries

To quantify BFP-BNIP3 intensities, Fiji v.1.5 was used. Briefly, maximum intensity projections of 21 z-stacks were generated (+-10 stacks from the middle z-stack rounded down). Background noise was subtracted from the ATP5a (mitochondria) and BNIP3 channels with rolling ball radius 30, and the mask of mitochondria was generated using ATP5a signal with default thresholding method. To obtain BNIP3 intensity on mitochondria, BNIP3 raw integrated pixel intensity within the mitochondrial mask was normalized to the mitochondrial mask surface area.

### *GSK3* and *BNIP3* mRNA quantification

For GSK3 RNAi efficacy quantification, females expressing maternal triple driver (MTD; COG-Gal4:VP16, nos-Gal4-VP16, Gal4-nos.NGT) and a UAS-shRNA targeting GSK3 were crossed to wild-type males and maintained on apple juice plates with yeast paste. After synchronization of egg lay, embryos were collected for two hours and dechorionated using liquid bleach. Total RNA was extracted from embryos using Tri-Reagent (BioShop Canada, Cat#TRI118), chloroform (Sigma-Aldrich, 472476), 2-propanol (Sigma-Aldrich, I9516) and ethyl alcohol (Commercial Alcohols, 22734). Purified RNA was treated with Turbo DNAse (ThermoFisher Scientific, 2238G2). For cDNA synthesis, 2000 ng of RNA was used with SuperScript II Reverse Transcriptase kit (ThermoFisher Scientific, 18064014) and oligo dT-20mer (Integrated DNA Technology, 51-01-15-01). Quantitative PCRs were carried out on 1/50 of reverse transcription reaction and 300 nM of each primer using the SensiFAST SYBR No-ROX kit (FroggaBio, BIO-98050) and a Bio-Rad CFX384/C1000 Touch system (Bio-Rad). The PCR program was as follows: 2 min at 95 °C; 45 cycles of 95 °C for 5 s and 60 °C for 30 s. GSK3 (Fwd: *5’*-*TTGGCGCCATCAATTATACA-3’* and Rev: *5’-TTGGATTCATTTCGCGTATC-3’*) expression results were normalized to the mean of value obtained of CG8187 (Fwd: *5’-AGACGCCTGGAAGTAAGCAG-3’* and Rev: *5’-GTAAGGCGGCTCAACTTGTC-3’*), CG2698 (Fwd: *5’-CTTCAGCATTTGTGGCAGAC-3’* and Rev: *5’-ATGTGTCGCTCTGGTGACTG-3’*) and Und (Fwd: *5’-GCAAGAAAAGCGGTCAGACT-3’* and Rev: *5’-CGTGTTGATACGGTCCAGAG-3’*). Gene knockdown was normalized to the relative expression level in LexA knockdown embryos. Results were calculated using the following formula: ΔΔCt = 2^-(ΔCtRNAi – ΔCtLexA RNAi), where ΔCt = Ct (gene) – Ct (mean of CG8187, CG2698 and Und).

mRNA quantification in BNIP3 mutant flies was done similarly except that whole flies were used and the primers used for BNIP3 were *5’-CAAGCGTGGCATACTCCTTA-3’* and *5’-GTTGGAATACGGTTGGCTGT-3’*.

### BNIP3 protein purification

#### Cloning

Wildtype and phospho-mutant forms of GST-BNIP3^ΔTM^ fusion constructs were generated by cloning the coding sequence of BNIP3 isoform A into the pGEX-4T1 (Addgene) vector for optimized expression in Escherichia coli. Coding sequences of BNIP3 and its mutant forms, excluding its transmembrane domain, were ordered as gene fragments from Twist Bioscience, PCR amplified [RP8.1] using 2x GB-AMP PaCeR HP Master Mix (GeneBio Systems) and cloned using Gibson assembly using NEBuilder HiFi DNA Assembly Master Mix (NEB E2621).

#### Expression

GST-BNIP3^ΔTM^ fusion constructs were purified from BL21 competent Escherichia coli cells. pGEX- GST-BNIP3^ΔTM^ plasmids were transformed into BL21 cells and transformants were selected on LB plates containing ampicillin (BioShop AMP201) and chloramphenicol. (BioShop CLR201) 20mL of 2YT media (0.5% NaCl, 1.6% Tryptone, 1% Yeast Extract w/v) containing ampicillin + chloramphenicol was inoculated with single colonies as biological replicates and grown overnight (∼16hrs) at 37°C in a shaking incubator. The following day, 500mL of 2YT media was inoculated with 5mL of the overnight culture and grown at 37°C in a shaking incubator until an O.D 600nm of 0.6-0.8 was reached. Protein expression was induced with 0.25mM IPTG and incubated at 18°C in a shaking incubator overnight (∼16hrs).

#### Purification

Bacterial cultures were pelleted at 4000 RPM for 30 mins at 4°C and stored at −80°C until ready for purification, for up to one week. Cell pellet was thawed and lysed in lysis buffer (1x BugBuster [Millipore Sigma 70921-4], 1M NaCl, 1mM AEBSF, 2mM benzamidine, 2ug/mL leupeptin, 2ug/mL pepstatin, 1mM DTT, 1% Sarkosyl) and incubated on ice for 30 mins. Cell lysate was then centrifuged at 15,000 g for 30 mins at 4°C. Supernatant from the lysate was added to glutathione agarose bead (Millipore Sigma G4510) slurry (1/1000 of bacterial culture volume), that was washed three times in wash buffer (150mM NaCl, 50mM Tris pH 7.4) to make a 1:1 bead to buffer slurry and incubated at 4°C for one hour. Beads were then pelleted from the lysate supernatant and loaded onto a Poly-Prep® Chromatography Column (Bio-Rad #7311550) and allowed to settle. The loaded column was washed six times with 1mL 1X PBST (1% Triton) each, and five times with 1X PBS. 10uL of the last wash fraction was tested with Bradford Protein Assay (Bio-Rad #5000006) to ensure no protein was washed out. Protein was then eluted with 0.8 column volume of elution buffer (250mM KCl, 100mM Tris pH 8.0, 30mM Glutathione [Millipore Sigma G4251]) into six fractions. Using Bradford Protein Assay, fractions with highest protein concentration were determined and pooled for dialysis. The pooled fractions were dialyzed using SnakeSkin^TM^ Dialysis Tubing bags (Thermo Scientific #88243) in dialysis buffer (20mM HEPES, 150mM KCl, 20% Glycerol), once overnight and once again for at least 4 hrs at 4°C. Dialyzed protein solutions were flash frozen in liquid nitrogen in 100uL aliquots and stored in −80°C after analyzing with SDS-PAGE and immunoblotting.

### *In vitro* BNIP3 phosphorylation

#### ADP-Glo Assay

The following assay was performed as outlined in the technical manual of ADP-Glo^TM^ Kinase Assay kit (Promega V6930) using commercially available human GSK3B (Sino Biological #10044-H07B). The optimal amount of kinase was determined to be 0.25µg per reaction using histone as a model substrate to find the linear range, and the optimal amount of BNIP3 substrate was determined to be ∼60-70ng, based on greatest observed difference in luminescence with and without the kinase (see technical manual). Thus, the subsequent kinase reactions were run using 0.25µg of hGSK3 and 70ng of purified BNIP3 in a total of 5µL reaction volume for 1 hr at room temperature, before adding the detection reagent. Luminescence was measured with an integration time of 1 second per well. All raw values (RLU) were normalized to internal negative controls, which are paired reactions that were run in the absence of the kinase.

#### Electromobility Shift Assay

Kinase reactions were performed using 0.25µg of hGSK3 and 10µg of BNIP3 substrate in alternate buffer (25mM HEPES pH 7.4, 20mM MgCl2, 1mM DTT, 0.5mM Ultrapure ATP) at 4°C for 72 hr, in a total reaction volume of 120µL. To inhibit GSK3, 2µM of CHIR99021 (Millipore Sigma SML1046) was added prior to incubation. To remove phospho-group, reaction mixture was treated with 400 units of λ-phosphatase (NEB P0753S), for 30 mins at 30°C, according to recommended protocol. 12µL of the reaction (containing ∼1µg of substrate) was run on SDS-PAGE gel for analysis by gel staining and immunoblotting with GST-tag probe, while a 10X diluted aliquot was run on SDS-PAGE gel for analysis by immunoblotting with His-tag probe. Gels were run on a 4-15% gradient gel (Bio-Rad # 4561086) at 150V for 1.5hrs. For staining, gels were incubated in InstantBlue® Coomassie Protein Stain (ISB1L) (Abcam ab119211) for 30 mins and destained overnight in MiliQ water. For immunoblotting, gels were transferred onto Immun-Blot PVDF Membrane (Bio-Rad #1620177) at 100V for 45 mins. Blots were probed with either anti-GST antibody (Millipore Sigma G7781, 1:2000) to detect the substrate or anti-His antibody (GenScript A00186S, 1:2000) to detect the kinase.

### Affinity purification mass spectrometry

#### Co-Immunoprecipitation

To generate BNIP3 constructs, BNIP3 CDSs with the necessary mutations were synthesized using Twist Bioscience, tagged N-terminally with FLAG, and cloned into pAc5.1 plasmid using NEBuilder HiFi DNA Assembly Master Mix (NEB E2621L). S2 cells were transfected with BNIP3 constructs using TransIT-Insect Reagent (Mirus Bio MIR 6105). Untransfected cells were used as a negative control. 2 days post transfection cells were harvested and lysed in lysis buffer (20mM Tris-HCl, 150mM NCl, 1mM EDTA, 1% NP40 containing 1x Roche protease inhibitor cocktail (Roche 11836170001)). Cell lysates were incubated with anti-FLAG beads (Thermo Scientific A36797) for 3hrs at 4°C with rotation. Samples were then processed for either SDS-PAGE followed by immunoblotting or for mass spectrometry.

#### Western blotting

Samples were run on freshly made 10% bis-acrylamide gels. Blots were stained with rabbit a-HA (CST, 3724S, 1:1000) or mouse anti-FLAG (Sigma, F1804-200UG, 1:5000) diluted in 1x PBSTB (1x PBS supplemented with 0.1% Tween-20 and 5% milk) followed by secondary antibodies (anti-Rabbit-HRP 1:8000 or anti-Mouse-HRP 1:8000).

#### Mass spectrometry

Cells were processed as described above except for TransIT-Insect Reagent (Mirus Bio MIR 6105), which was used for transfection. Protein concentration was determined using BCA assay. IPs were performed on 10 mg of protein lysate at a concentration of 10 mg/ml. IP’ed samples were resuspended in Tris-HCl (pH 8.0) and submitted to the NBCC proteomics at the Lunenfeld-Tanenbaum Research Institute. For data-dependent acquisition (DDA) LC-MS/MS, One-sixteenth of digested peptides were analyzed using a nano-HPLC (High-performance liquid chromatography) coupled to MS. One-sixteenth of the sample was loaded onto Evotip Pure per manufacturer instructions. Peptides were eluted from the Performance column (cat#: EV-1109, 8cm x 150µm with 1.5µm beads), heated at 40°C) with the 60SPD pre-formed acetonitrile gradient generated by an Evosep One system, and analyzed on a timsTOF Pro 2. The Evosep was coupled to timsTOF Pro 2 using a 20µm diameter emitter tip. The column toaster was set to 40°C. The total DDA protocol was 22 minutes. The MS1 scan had a mass range of 100-1700Da in PASEF mode. TIMS settings were accumulation and ramp time of 100ms (with 4 PASEF ramps and active exclusion at 0.4min), and within the mobility range (1/K0) of 0.85 to 1.3V·s/cm2. This was at a cycle time of 0.53s. The target intensity was set to 17,500 and intensity threshold set to 1750. 1+ ions were excluded from the fragmentation using a polygonal filter. The auto calibration was off. The raw files were searched using FragPipe v22.0 with MSFragger v4.1, IonQuant v1.10.27, and diaTracer v.1.3.

### Phospho-proteomic mass spectrometry analysis

Phospho-proteomic mass spectrometry experiments were performed by the SPARC BioCentre (The Hospital for Sick Children, Toronto, Ontario). Briefly, liquid chromatography with tandem mass spectrometry (LC-MS/MS) was performed using SDS-PAGE gel slices of protein samples stained with InstantBlue (Abcam). Briefly, gel bands were split into two for separate digestion with trypsin (ThermoScientific Pierce) and thermolysin (Promega), respectively. Reduction was performed using 10 mM DTT (60°C, 1 h) followed by alkylation using 55 mM iodoacetamide (25°C, 20 min in dark). Trypsin digestion (approximately 600 ng per sample, or approximately 1:50 to 1:100 protease:protein) was carried out at 37°C overnight, whereas thermolysin digestion (approximately 600 ng per sample) was carried out at 70°C for 1 h followed by incubation at 37°C overnight. Peptide extraction was performed using 5% formic acid in acetonitrile, where trypsin and thermolysin digests were combined and lyophilized before resuspension in 1% formic acid. LC-MS/MS was performed using an EASY-nLC 1200 chromatography system (ThermoFisher Scientific), coupled to an Orbitrap Fusion Lumos Tribrid mass spectrometer (ThermoFisher Scientific), using a gradient of 0.1% formic acid with 0.1% formic acid and 80% acetonitrile. The mass spectrometry data is provided in Table S2. Data analysis was performed using the software Scaffold 5 and Scaffold PTM (Proteome Software). The intact mass of unphosphorylated and phosphorylated samples was determined by liquid chromatography coupled to electrospray ionization mass spectrometry (LC-ESI-MS) using protein samples (40 μM) in SEC buffer (25 mM sodium phosphate pH 7.4, 100 mM NaCl, 0.1 mM EDTA).

### Cell cultures

Drosophila S2 cells were cultured in Express Five serum free medium (Gibco, 10486025) supplemented with 20mM L-glutamine (Sigma-Aldrich, G7513) and 1% penicillin-streptomycin (Sigma-Aldrich, P4333) and incubated at 25°C.

HeLa cells were cultured in DMEM (Sigma-Aldrich, D5796) supplemented with 10% FBS (Sigma-Aldrich, F1051) and 1% penicillin-streptomycin (Sigma-Aldrich, P4333) at 37°C, 5%CO2. For drug treatments, HeLa cells were transfected with a plasmid expressing mito-QC using Lipofectamine 3000 Transfection Reagent (Thermo Scientific, L3000008), allowed to recover overnight, and treated with 1 mM DFP and 5µM CHIR-99021 for 30 hours. For negative controls cells were treated with water and DMSO.

### Mitophagy quantification in cells

For imaging, cells were cultured and fixed directly in 8-well Nunc™ Lab-Tek™ II Chambered Coverglass (Thermo Scientific, 155409), washed with PBS, and fixed with 4% PFA diluted in PBS for 20 minutes. Cells were imaged in PBS on a Leica SP8 inverted scanning confocal microscope using a 63X, NA 1.4 immersion oil objective. Fields of views were chosen based on the brightfield channel. Mitophagy was quantified using Imaris v.9. Surfaces for mitochondria were generated from the GFP signal and the surfaces for mitochondria + mitolysosomes were generated using mCherry signal. Background was subtracted using the local contrast thresholding method with surface grain size = 0.1µm and the diameter of largest sphere = 1.35µm. Identical manual brightness threshold was used for GFP and mCherry and was kept the same throughout the experiment. Only the surfaces larger than 10.0 voxels were kept. To obtain mitophagy volume, the sum of volume of surfaces generated with GFP was subtracted from the sum of volume of surfaces generated with mCherry and normalized to the number of cells transfected.

### Statistical analysis

Statistical analysis was done using GraphPad Prizm v.10.2.0 for all data except for the mass spectrometry. For the IP-MS, significantly enriched interactors in BNIP3^WT^, BNIP3^pNull^ ^All^ and BNIP3^pMim^ ^All^ over negative control were identified using SAINT (Choi *et al*, 2011).

## SUPPLEMENTARY FIGURES

**Figure S1.**
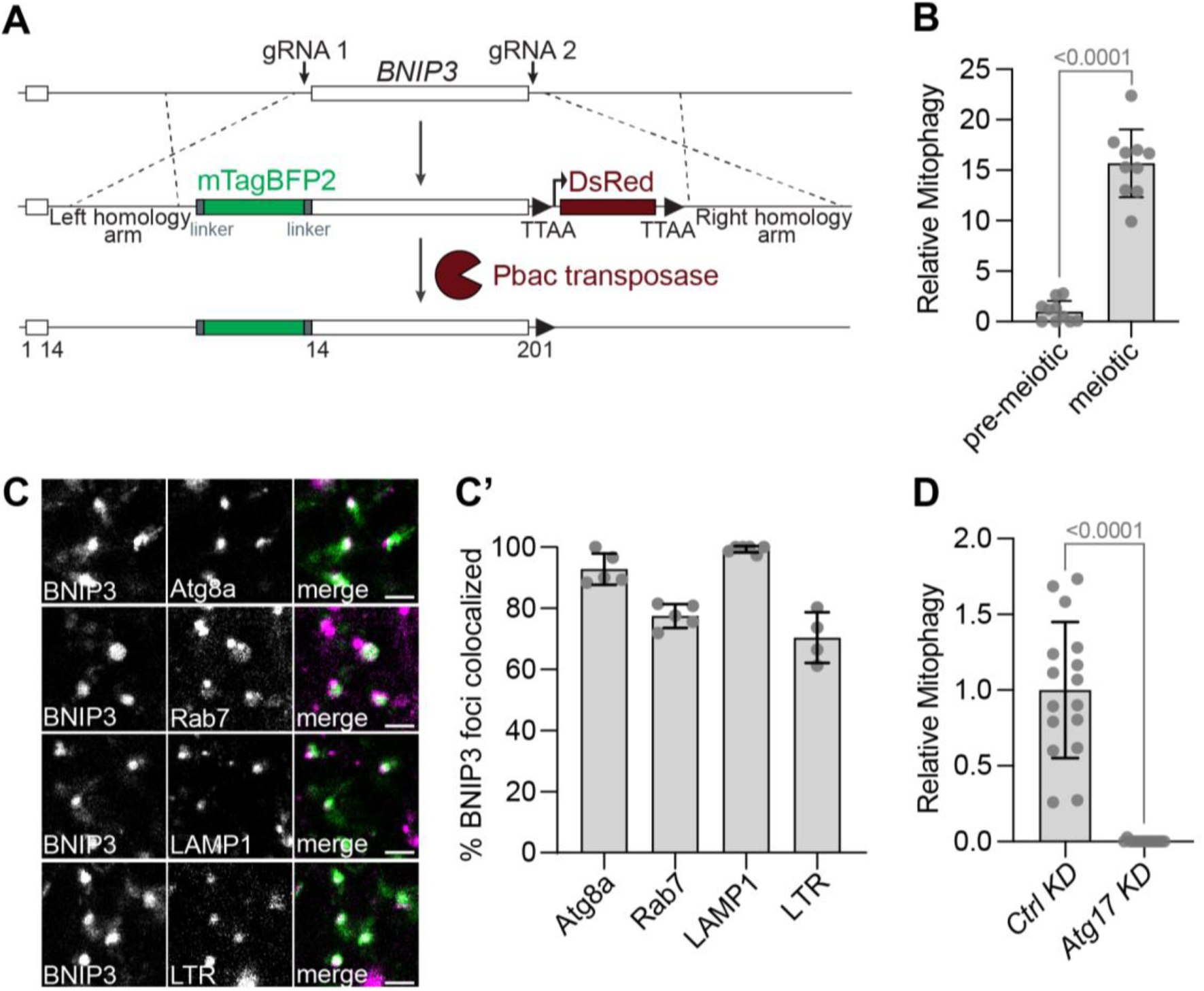
BNIP3-marked structures associate with autophagic compartments and require Atg17. **(A)** Scarless CRISPR/Cas9 scheme for generating endogenously BFP-tagged BNIP3 strains. Numbers indicate amino acid positions in BNIP3. **(B)** BNIP3-mediated mitophagy in pre-meiotic (region 1) and meiotic (region 2) germ cells quantified by overlap with Atg8a and normalized to the pre-meiotic samples. Values represent means ± s.d.; *p*-value, Mann-Whitney test. Genotype: *NGT40/UAS-lexA shRNA; BFP::BNIP3, 3xmCherry::Atg8a/+*. **(C)** Confocal micrographs showing BNIP3-positive structures (BNIP3, green) co-localizing with markers of the autophagic pathway (magenta), including mCherry-Atg8a, eYFP-Rab7, LAMP1-mCherry, and LysoTracker Red. Scale bars, 2.5 µm. Genotypes: *BNIP3 + Atg8a* = *BFP::BNIP3/3xmCherry::Atg8a | BNIP3 + Rab7 = BFP::BNIP3/eYFP^MYC^::Rab7 | BNIP3 + LAMP1 = BFP::BNIP3/LAMP1::3xmCherry | BNIP3 +* LTR *= BFP::BNIP3/+*. **(C’**) Quantification of the fraction of BNIP3 foci co-localizing with the indicated markers. Values represent means ± s.d. **(D)** Germline knockdown (*nos*-Gal4) of the Atg1 complex component *Atg17* disrupts BNIP3-dependent mitophagy, assessed by reduced overlap with Atg8a. Data are normalized to control (*LexA)* knockdown and shown as means ± s.d.; *p*-value, Mann-Whitney test. Genotypes: *Ctrl KD* = *NGT40/+; BFP::BNIP3, 3xmCherry::Atg8a/UAS-LexA shRNA | Atg17 KD* = *NGT40/+; BFP::BNIP3, 3xmCherry::Atg8a/UAS-Atg17 shRNA*.

**Figure S2.**
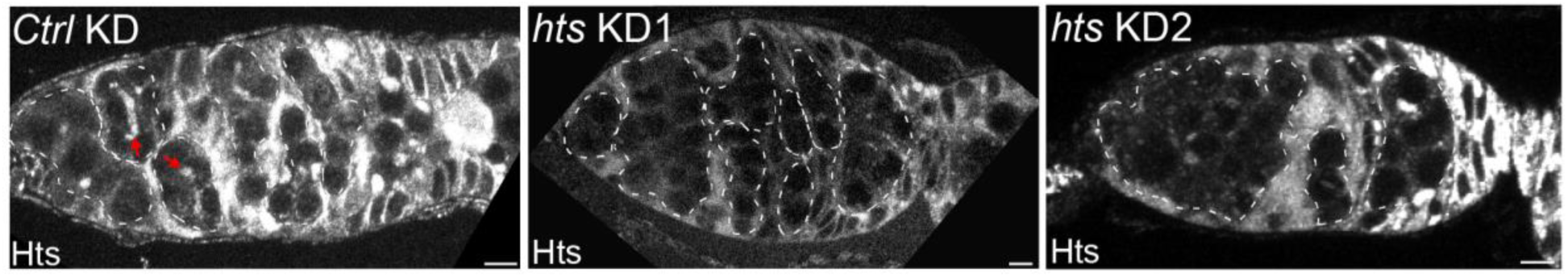
*hts* knockdown effectively reduces Hts protein and impairs BNIP3 mitophagy. Immunofluorescence confocal micrographs of control and *Hts* germline (*nos*-Gal4) knockdown ovaries stained with anti-Hts to mark fusomes and somatic tissues. Max intensity projections are shown. Red arrows mark the fusome. White dashed lines outline the germline. Scale bars, 5 µm. Genotypes: *Ctrl KD* = *NGT40/+; UAS-Lex shRNA/+ | Hts KD1* = *NGT40/+; UAS-hts shRNA 1/+ | Hts KD2* = *NGT40/+; UAS-hts shRNA 2/+*.

**Figure S3.**
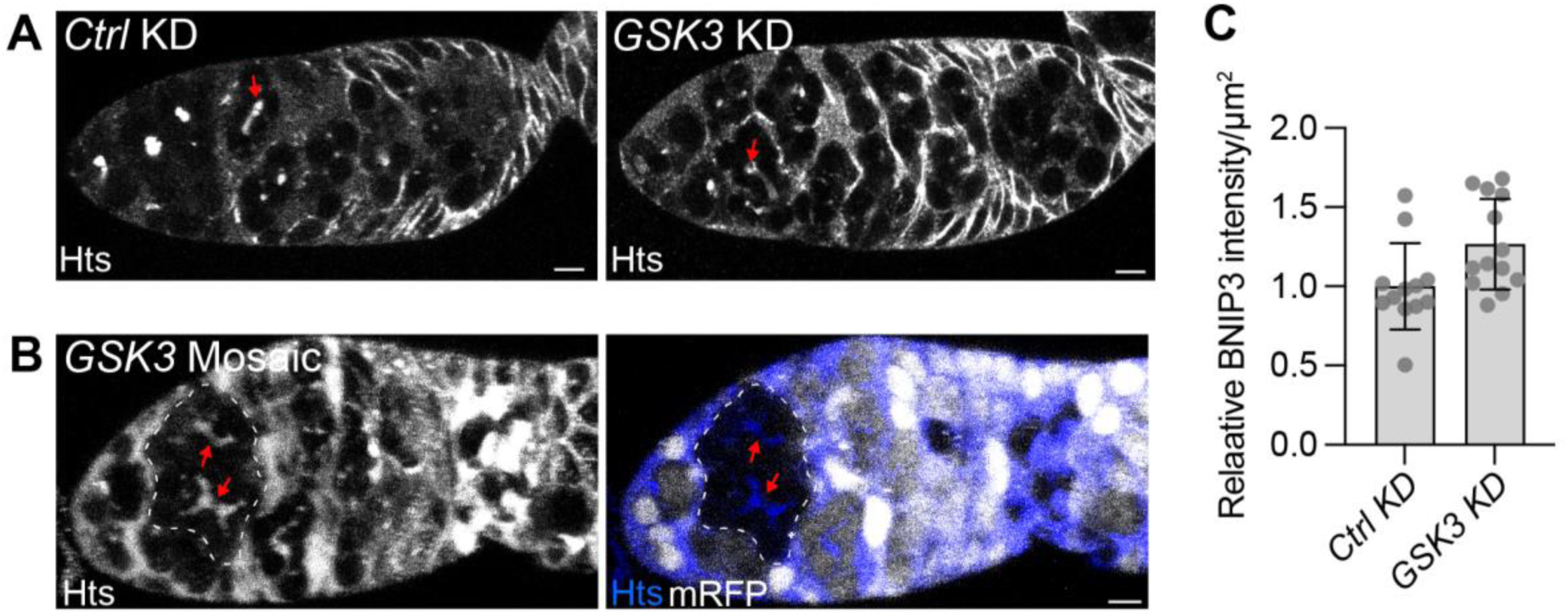
GSK3 is dispensable for fusome formation and maintenance. **(A)** Immunofluorescence confocal micrographs of control and *GSK3* germline (*nos*-Gal4) knockdown ovaries stained with anti-Hts to mark fusome (red arrows) and somatic tissues. Max intensity projections are shown. Scale bar, 5 µm. Genotypes: *Ctrl KD* = *NGT40/+; UAS-LexA shRNA/+ | GSK3 KD* = *NGT40/+; UAS-GSK3 shRNA/+*. **(B)** Immunofluorescence confocal micrograph of *GSK3* mutant mosaic ovaries with *GSK3* mutant cells marked by loss of mRFP (dashed outline). Fusomes (red arrows) and somatic tissues are visualized by anti-Hts staining. Max intensity projections are shown. Scale bar, 5 µm. Genotype: *Ubi*-*mRFP, hsFLP, FRT19A/GSK3^1^, FRT19A;;BFP::BNIP3/+*. **(C)** Quantification of BFP-BNIP3 protein levels in control and *GSK3* germline (*nos*-Gal4) knockdown ovaries. Values are normalized to *ctrl KD* and shown as means ± s.d. Genotypes: *Ctrl KD* = *NGT40/+; UAS-LexA shRNA/BFP::BNIP3, 3xmCherry::Atg8a | GSK3 KD = NGT40/+; UAS-GSK3 shRNA/BFP::BNIP3, 3xmCherry::Atg8a*.

**Figure S4.**
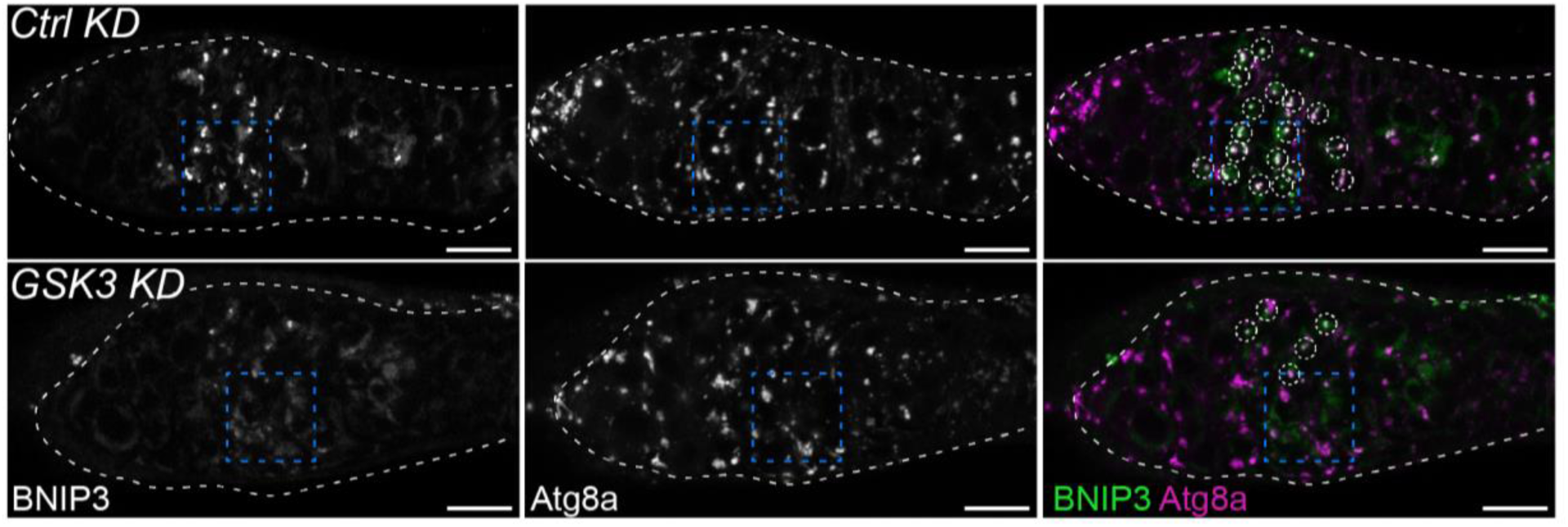
GSK3 as required for BNIP3-mediated germline mitophagy. Confocal micrographs of BFP-BNIP3 and mCherry-Atg8a in control and *GSK3* germline (*nos*-Gal4) knockdown ovaries. Max intensity projections are shown. White dashed lines outline the ovary. White dashed circles outline BNIP3- and Atg8a-positive foci. Blue dashed boxes indicate zoomed region presented in Fig. 2. Scale bars, 10 µm. Genotypes: *Ctrl KD* = *NGT40/+; BFP::BNIP3, 3xmCherry::Atg8a /UAS-Lex shRNA | GSK3 KD* = *NGT40/+; BFP::BNIP3, 3xmCherry::Atg8a/UAS-GSK3 shRNA*.

**Figure S5.**
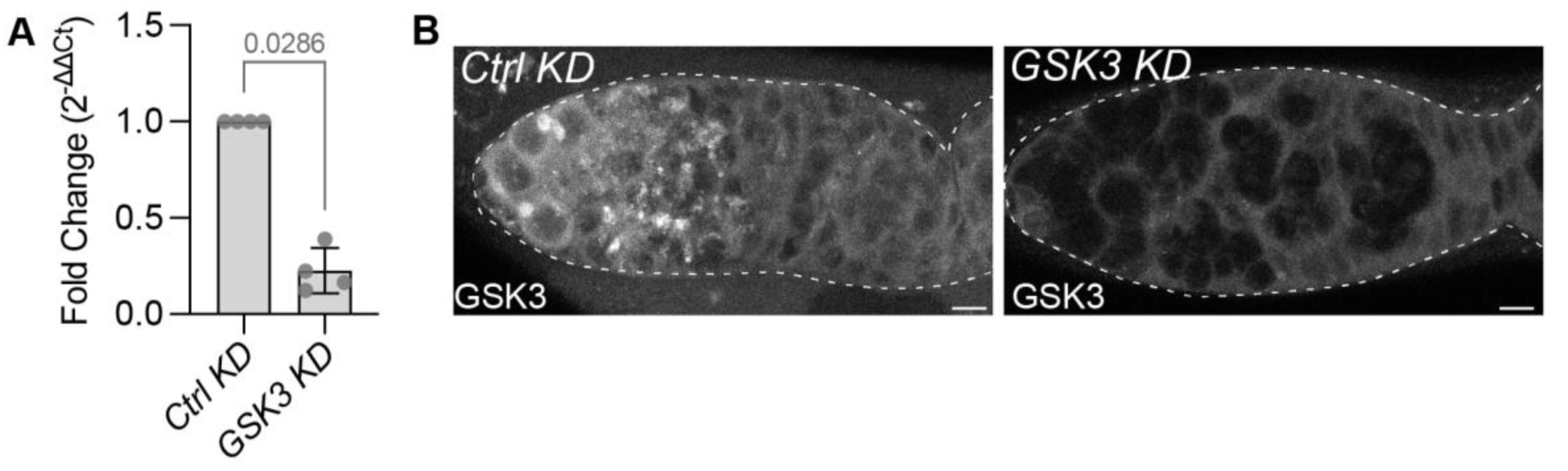
Germline knockdown of *GSK3* effectively silences GSK3. **(A)** *GSK3* mRNA levels in control and *GSK3* germline (*matα*-Gal4) knockdown samples. RT-qPCR was conducted on eggs from females expressing either control or *GSK3* RNAi. Values are normalized to control eggs and represent means ± s.d. *p*-value, Mann-Whitney test. Genotypes: *Ctrl KD* = *GSK3::GFP/+; matα*-Gal4*/+; matα*-Gal4/*UAS-LexA shRNA | GSK3 KD* = *GSK3::GFP/+; matα*-Gal4*/+; matα*-Gal4/*UAS-GSK3 shRNA*. **(B)** Confocal micrographs of GSK3-GFP ovaries with control or *GSK3* knocked down in the germline (*nos*-Gal4). Max intensity projections are shown. White dashed lines outline the ovary. Scale bars, 5 µm. Genotypes: *Ctrl KD* = *GSK3::GFP/+; NGT40/+;UAS-LexA shRNA/+* | *GSK3 KD* = *GSK3::GFP/+; NGT40/+; UAS-GSK3 shRNA/+*.

**Figure S6.**
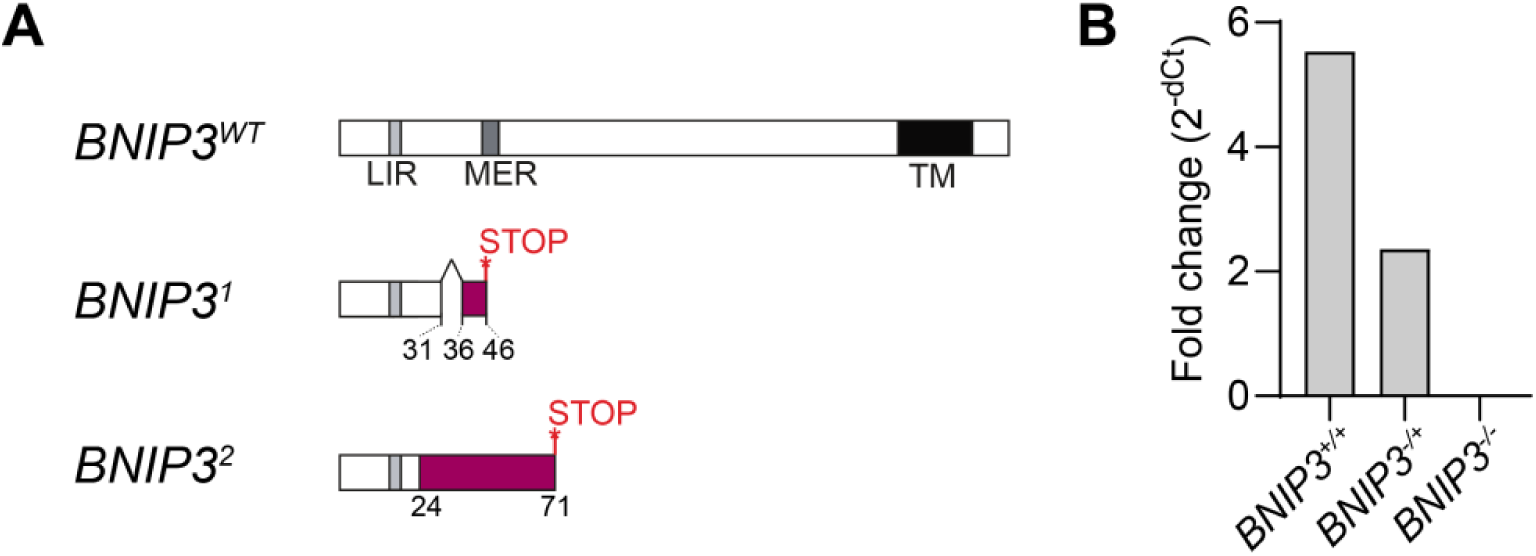
*BNIP3^1/2^* transheterozygotes exhibit loss of *BNIP3* transcripts. **(A)** A cartoon showing the *BNIP3* alleles generated with CRISPR/Cas9 genome editing. Purple colour shows frame-shifted amino acid sequences generated due to indels. **(B)** *BNIP3* mRNA levels in wildtype, heterozygous, and transheterozygous *BNIP3* flies. RT-qPCR was conducted on whole flies. Values are normalized to the average of housekeeping genes. Genotypes: *BNIP3^+/+^ = +/TM3 | BNIP3^-/+^ = BNIP3^2^/TM6B | BNIP3^-/-^ = BNIP3^1^/BNIP3^2^*.

**Figure S7.**
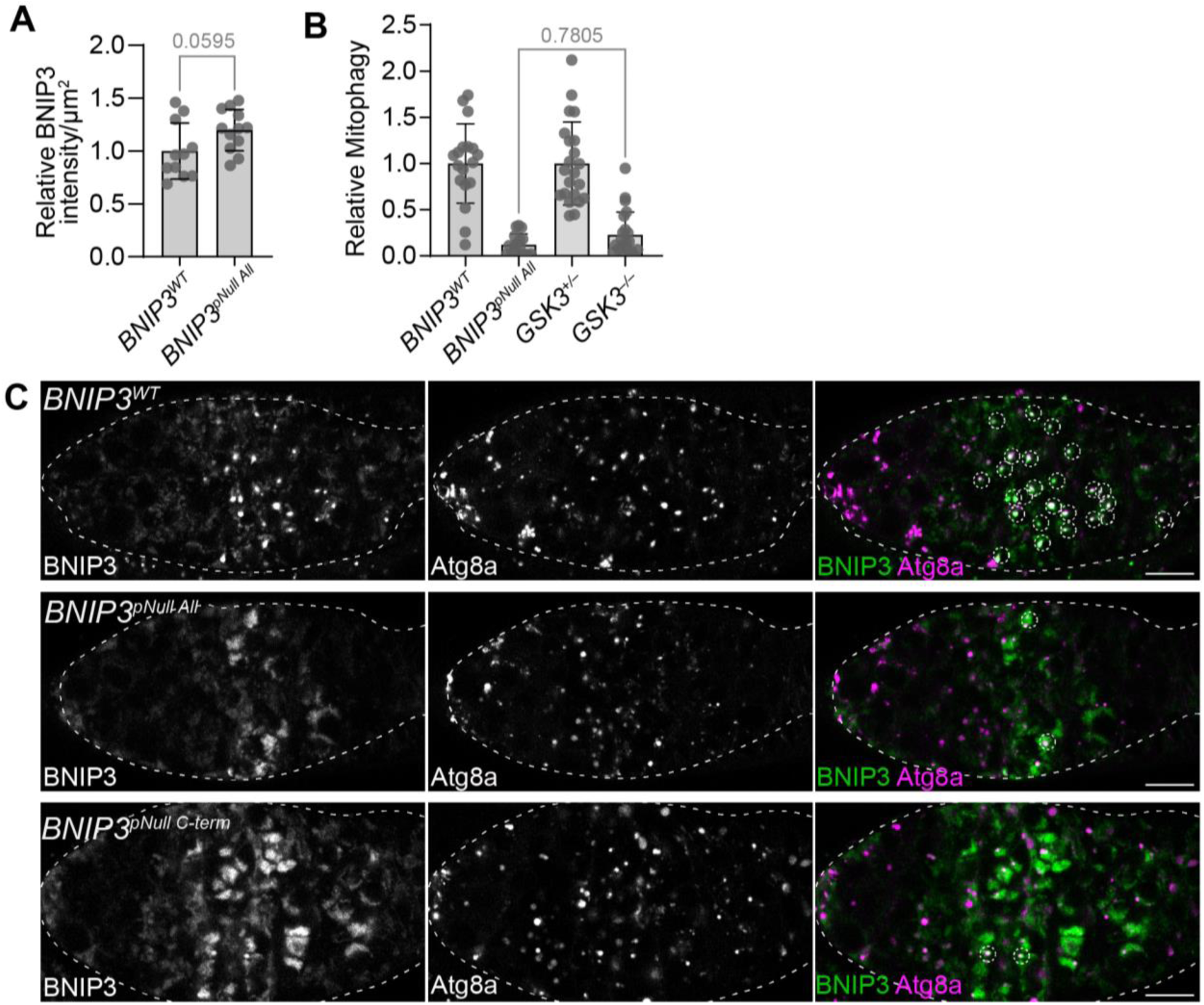
BNIP3 phosphorylation sites are essential for its activity. **(A)** Quantification of BFP-BNIP3 protein levels in wild-type or phospho-null *BNIP3* flies. Values are normalized to *BNIP3^WT^* and shown as means ± s.d. *p*-values, Mann–Whitney test. Genotypes: *BNIP3^WT^* = *3xmCherry::Atg8a/+; BFP-BNIP3^WT^/BNIP3^−^ | BNIP3^pNull^ ^All^*= *3xmCherry::Atg8a/+; BFP-BNIP3^10S/T>A^/BNIP3^−^*. **(B)** Quantification of mitophagy in wild-type or phospho-null *BNIP3* compared with heterozygous or homozygous *GSK3* mutant cells. Values are normalized to respective controls and shown as means ± s.d. *p*-values were calculated using one-way ANOVA with Tukey’s correction. Only the *p*-value between the phospho-null and *GSK3* mutant is indicated. *BNIP3^WT^*and *BNIP3^WT^ and BNIP3^pNull^ ^All^* data are from Fig. 4B. *GSK3^+/–^* and *GSK3^−/−^* data are from Fig. 2B’. Genotypes: *BNIP3^WT^* = *3xmCherry::Atg8a/+; BFP::BNIP3^WT^/BNIP3^−^ | BNIP3^pNull^ ^All^* = *3xmCherry::Atg8a/+; BFP::BNIP3^10S/T>A^/BNIP3^−^ | GSK3* mosaics = *Ubi-mRFP, hsFLP, FRT19A/ GSK3^1^, FRT19A;;BFP::BNIP3/+*. **(C)** Confocal micrographs of BFP-BNIP3 in ovaries from flies expressing wildtype and phospho-dead *BNIP3.* Max intensity projections are shown. Quantifications shown in Fig. 4B. White dashed lines outline ovaries. White dashed circles outline BNIP3- and Atg8a-positive foci. Scale bars, 10 µm. Genotypes: *BNIP3^WT^* = *3xmCherry::Atg8a/+; BFP::BNIP3^WT^/BNIP3^−^ | BNIP3^pNul^ ^All^* = *3xmCherry::Atg8a/+; BFP::BNIP3^10S/T>A^/BNIP3^−^ | BNIP3^pNull^ ^C-term^* = *3xmCherry::Atg8a/+; BFP::BNIP3^8S/T>A^/BNIP3^−^*.

**Figure S8.**
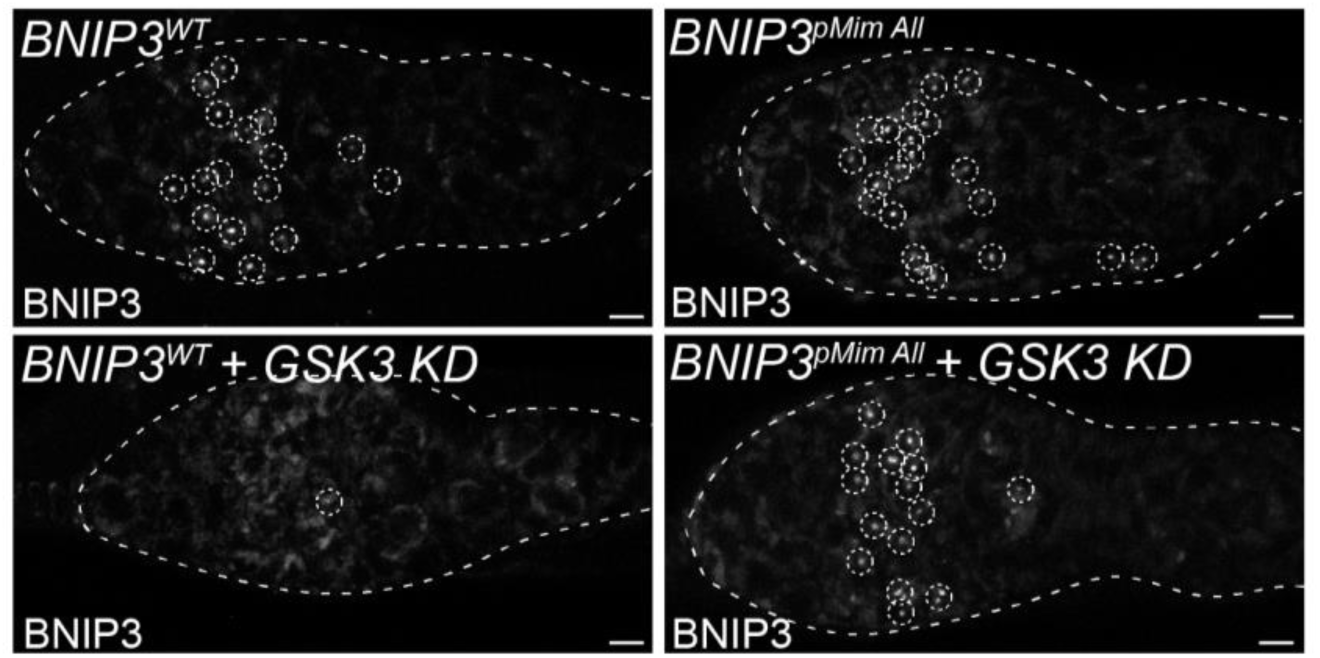
GSK3 acts upstream of BNIP3 to promote its phosphorylation and activation. Confocal micrographs of BFP-BNIP3 in ovaries expressing wildtype or phospho-mimetic (10S/T>D) *BNIP3* with or without *GSK3* germline (*nos*-Gal4) knockdown. Max intensity projections are shown. Quantification shown in Fig. 4C. White dashed lines outline ovaries. White dashed circles outline BNIP3 foci. Scale bars, 5 µm. Genotypes: *BNIP3^WT^*= *BFP::BNIP3^WT^; BNIP3^−^/BNIP3^−^ | BNIP3^pMim^ ^All^* = *BFP::BNIP3^10S/T>D^; BNIP3^−^/BNIP3^−^ | BNIP3^WT^ + GSK3 KD* = *NGT40/BFP::BNIP3^WT^; BNIP3^−^/BNIP3^−^, UAS-GSK3 shRNA | BNIP3^pMim^ ^All^ + GSK3 KD* = *NGT40/BFP::BNIP3^10S/T>D^; BNIP3^−^/BNIP3^−^, UAS-GSK3 shRNA*.

**Figure S9.**
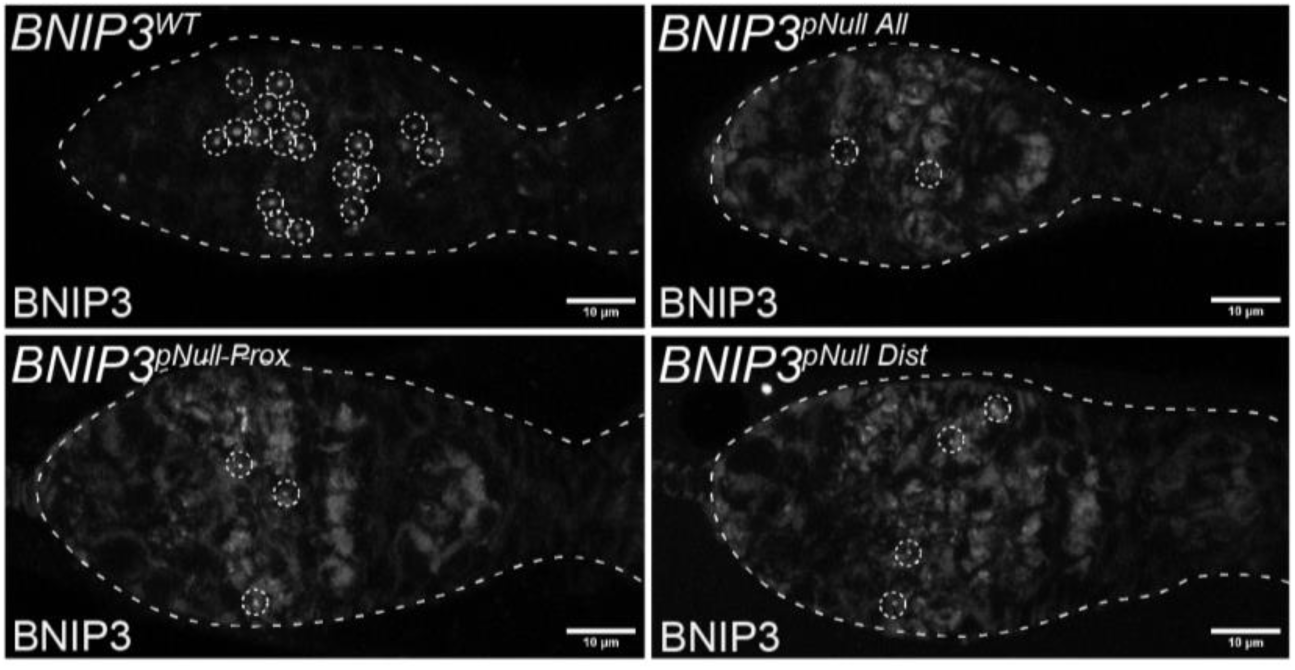
*BNIP3* phosphorylation sites are required for mitophagy. Confocal micrographs of BFP-BNIP3 in ovaries expressing wildtype or phospho-null *BNIP3* mutants (All, Prox, and Dist). Max intensity projections are shown. Quantification shown in Fig. 4D. White dashed lines outline ovaries. White dashed circles outline BNIP3 foci. Scale bars, 10 µm. Genotypes: *BNIP3^WT^* = *BFP::BNIP3^WT^; BNIP3^−^/BNIP3^−^ | BNIP3^pNull^ ^All^* = *BFP::BNIP3^10S/T>A^; BNIP3^−^/BNIP3^−^ | BNIP3^pNull^ ^Prox^* = *BFP::BNIP3^3S/T>A^; BNIP3^−^/BNIP3^−^ | BNIP3^pNull^ ^Dist^* = *BFP::BNIP3^5S/T>D^; BNIP3^−^ /BNIP3^−^*.

**Figure S10.**
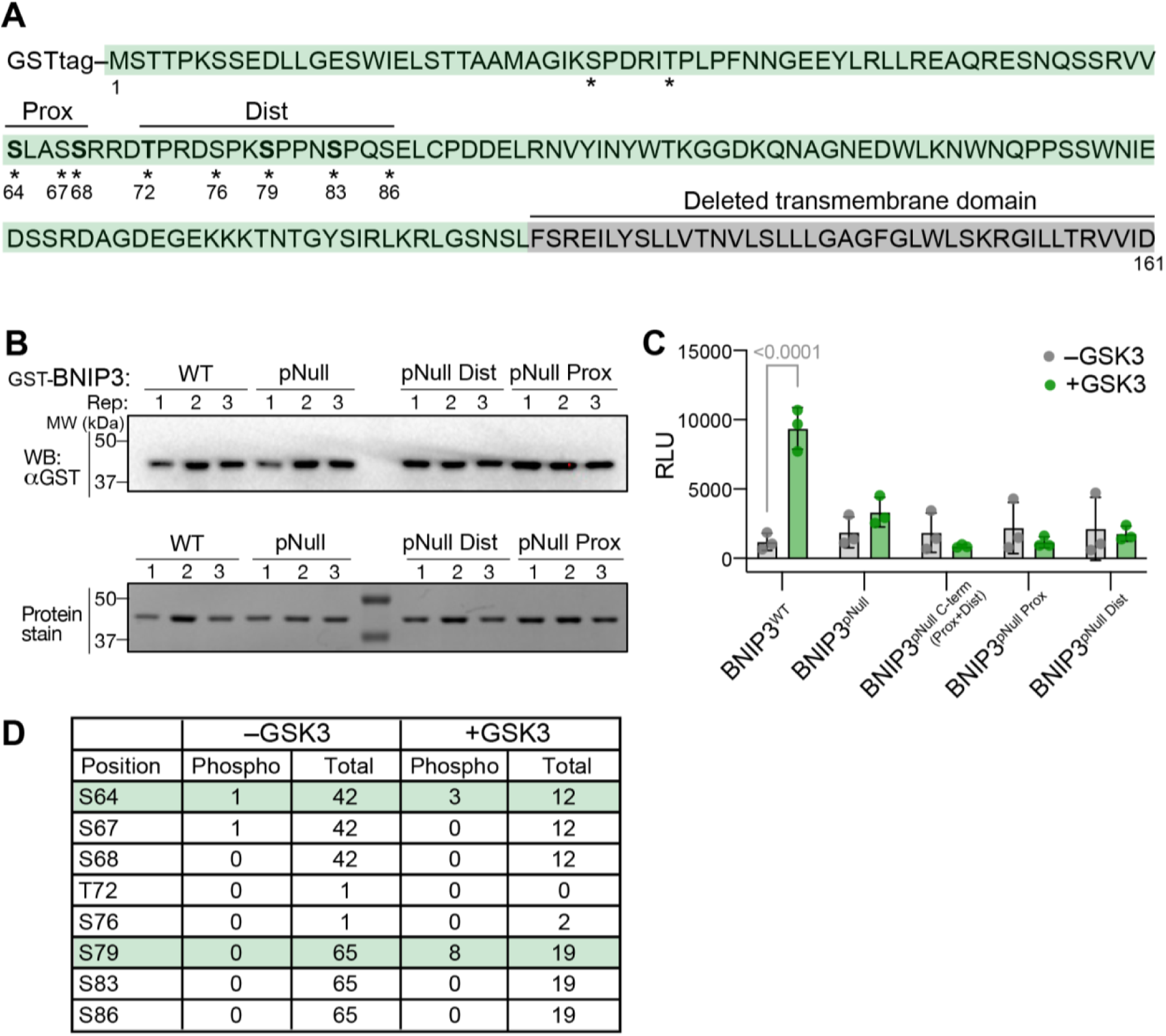
Purification and phosphorylation analysis of BNIP3. **(A)** Schematic showing the protein sequence of GST-tagged BNIP3 (green) with its transmembrane domain deleted (grey highlight) purified from *E. coli*. Mutated candidate phosphorylation sites are indicated with stars, and residues that fit the GSK3 consensus motif are bolded. **(B)** Purified GST-BNIP3 and its phospho-mutant variants visualized using western blotting with anti-GST, and InstantBlue Coomassie stained acrylamide gels. Three replicates shown per condition. **(C)** Quantification of *in vitro* kinase activity as measured by relative luminescence unit (RLU) using ADP-Glo kit. Plotted are raw RLU values measured in the presence and absence of the kinase (paired for each substrate). Each dot represents an independent experimental mean of technical triplicates. Normalized data shown in Fig. 5A. *p*-values, 2way ANOVA with Šídák’s multiple comparisons test. Only significant *p*-values (<0.05) are shown. **(D)** Table summarizing spectra counts from a LC-MS/MS experiment. BNIP3 was incubated with and without hGSKB and peptides were analyzed by mass spectrometry.

**Figure S11.**
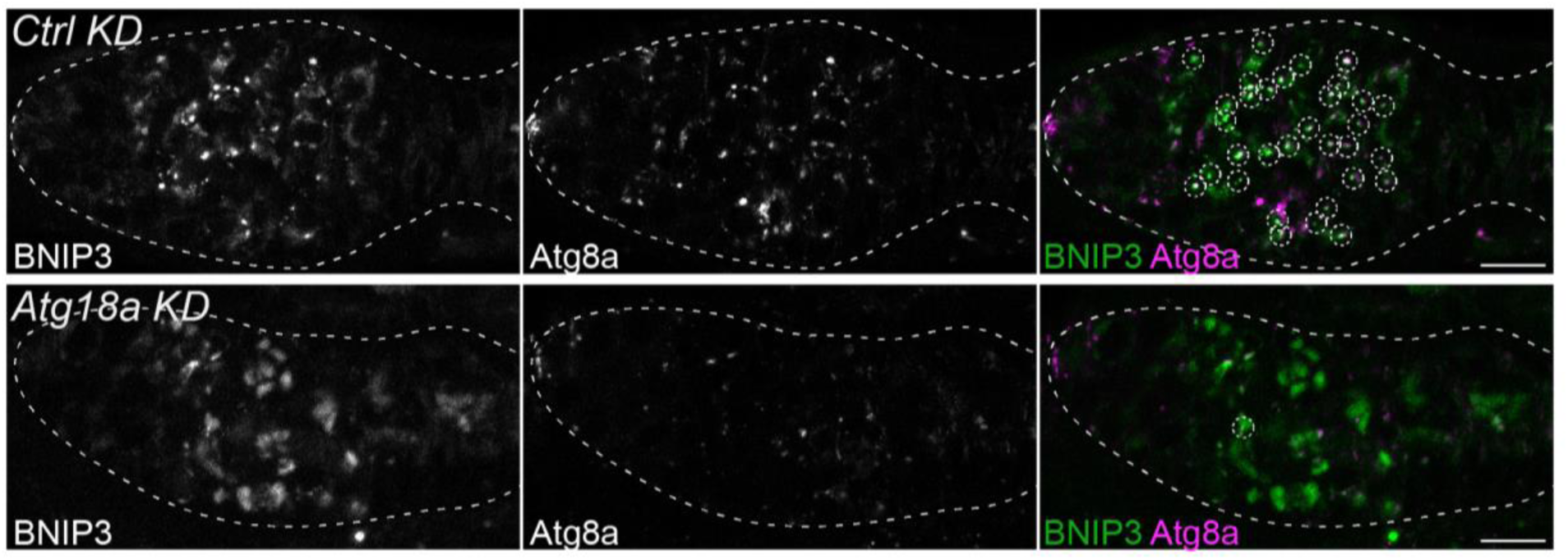
The knockdown of *Atg18a* impairs mitophagy in the *Drosophila* germline. Confocal micrographs of BFP-BNIP3 and mCherry-Atg8a in control and *Atg18a* germline (*nos*-Gal4) knockdown ovaries. Max intensity projections are shown. Quantification shown in Fig. 6D. White dashed lines outline ovaries. White dashed circles outline BNIP3- and Atg8a-positive foci. Scale bar, 10µm. Genotypes are the same as in Fig. 6D

**Figure S12.**
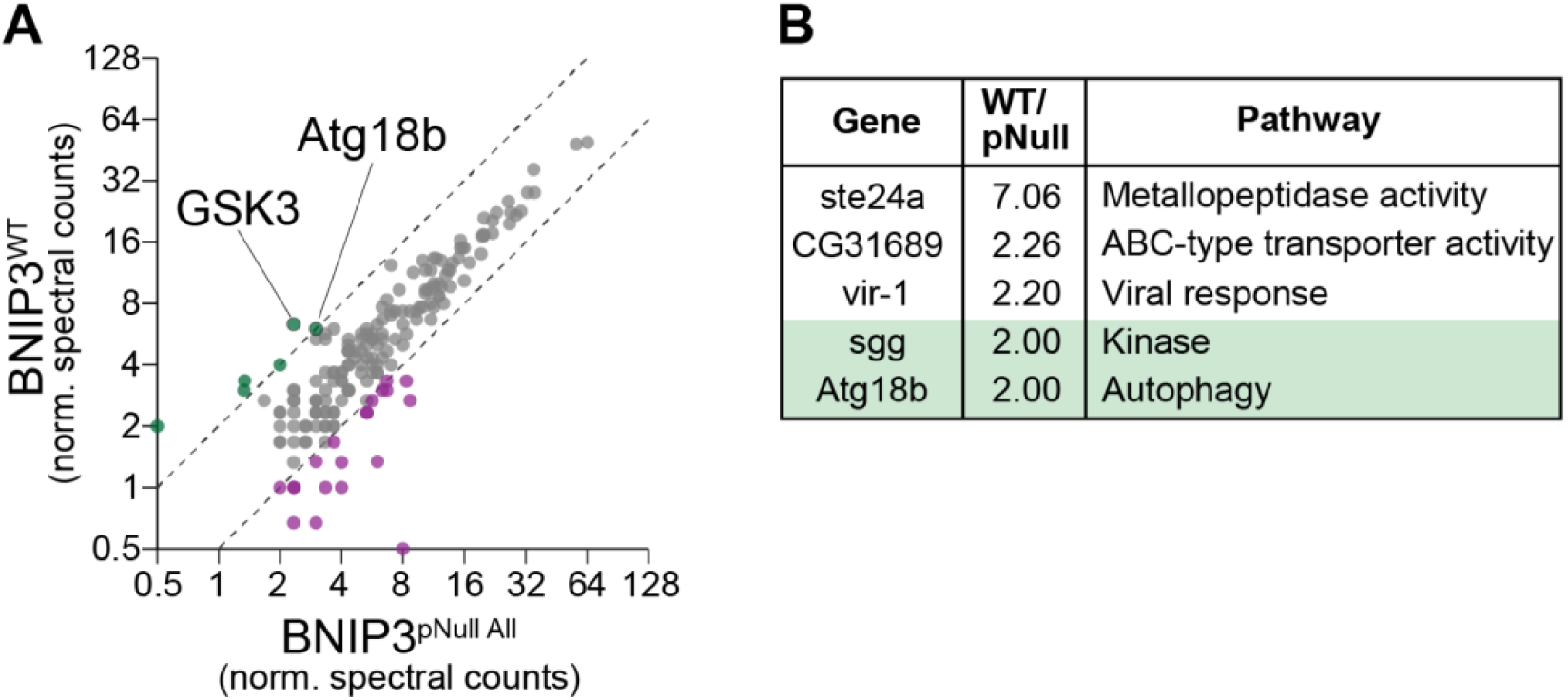
WT BNIP3 interacts with Atg18b and GSK3. **(A)** A scatterplot showing the normalized spectral counts in wildtype BNIP3 (BNIP3^WT^) and phosho-null BNIP3 (BNIP3^pNull^ ^All^). Spectral counts are averaged across three replicates. Dashed lines indicate 2-fold difference. Shown are interactors significantly enriched in the pulldowns relative to the negative control with BFDR ≤ 0.05. **(B)** Interactors enriched 2-fold in WT BNIP3 over phospho-dead BNIP3.

**Figure S13.**
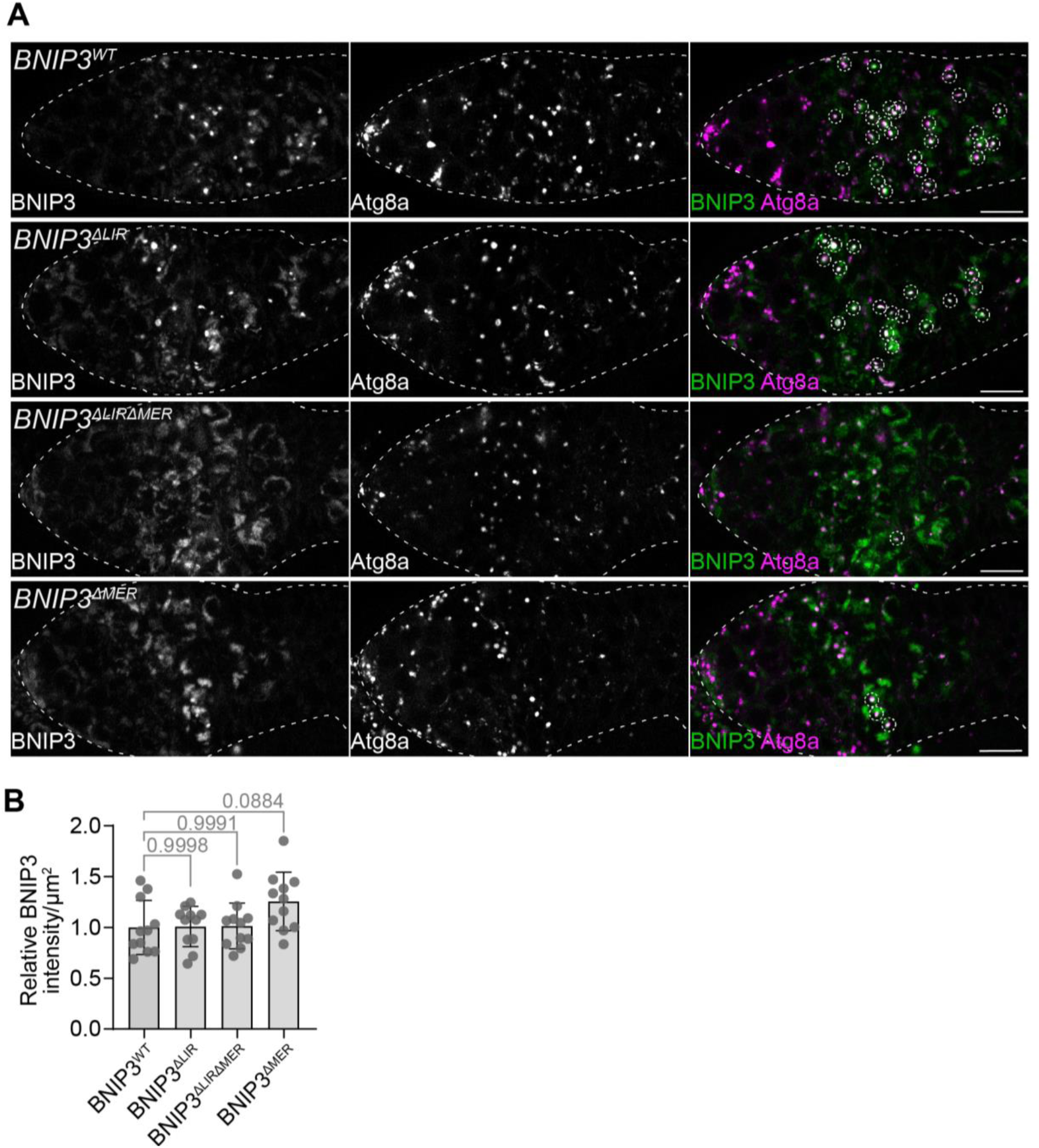
BNIP3 MER is required for mitophagy. **(A)** Confocal micrographs of BFP-BNIP3 in ovaries expressing wildtype or *BNIP3* mutants lacking the LIR, MER or both. Max intensity projections are shown. Quantification shown in Fig. 6F. White dashed lines outline ovaries. White dashed circles outline BNIP3- and Atg8a-positive foci. Scale bars, 10 µm. Genotypes: *BNIP3^WT^* = *3xmCherry::Atg8a/+; BFP::BNIP3^WT^/BNIP3^−^ | BNIP3^ΔLIR^* = *3xmCherry::Atg8a/+; BFP::BNIP3^ΔLIR^/BNIP3^−^ | BNIP3^ΔLIRΔMER^* = *3xmCherry::Atg8a/+; BFP::BNIP3^ΔLIRΔMER^/BNIP3^−^ | BNIP3^ΔMER^* = *3xmCherry::Atg8a/+; BFP::BNIP3^ΔMER^/BNIP3^−^*. **(B)** Quantification of BFP-BNIP3 protein levels from (A). Values are normalized to BNIP3^WT^ and shown as mean ± s.d. *p*-values, one-way ANOVA with Tukey’s correction. Only *p*-values between the wildtype and the mutants are shown.

**Table S1.** List of Drosophila strains used.

**Table S2.** All peptides detected by phospho-proteomic mass spectrometry of BNIP3 (lacking the transmembrane domain) incubated with or without hGSK3B.

**Table S3.** Interactors enriched ≥2-fold in phospho-mimetic BNIP3 immunoprecipitations relative to phospho-mutant BNIP3 immunoprecipitations (BFDR ≤ 0.01). Only interactors whose transcripts are enriched in meiotic germ cells are shown. Interactors are ordered by meiotic transcript enrichment.

**Table S4.** Interactors enriched ≥2-fold in wildtype BNIP3 immunoprecipitations relative to phospho-mutant BNIP3 immunoprecipitations (BFDR ≤ 0.01).

**Table S5.** Sequences of BNIP3 structure-function mutants generated with CRISPR.

